# Does early-life food shortage alter the effect of elevated temperature on female life history?

**DOI:** 10.1101/2024.07.24.605022

**Authors:** Meng-Han Joseph Chung, Chenke Zang, Diego Moura-Campos, Michael D. Jennions, Megan L. Head

## Abstract

1. Global warming is reducing prey availability in many aquatic systems, raising questions about the combined effects of higher temperatures and lower food availability on fish life histories and reproductive output.
2. In ectotherms, higher temperatures accelerate growth and promote an earlier onset of reproduction. However, when fish have less food during development, resource depletion might constrain these temperature-driven processes.
3. We manipulated water temperature (24 or 28°C) and early-life food availability (control or restricted) for female guppies (*Poecilia reticulata*). We measured how both factors affected key life history traits (growth, reproduction, survival, self-maintenance).
4. Higher temperature significantly affected female life histories. Females at 28°C matured at a larger size, but then grew more slowly and produced fewer, smaller offspring than females at 24°C. The effect of temperature on reproduction persisted even after controlling for body size, suggesting there was a shift in the fecundity-size relationship.
5. Adult mortality was greater at 28°C. Higher temperature also resulted in a longer gut, potentially enhancing resource acquisition, but a higher temperature did not affect immunity or telomere length of the surviving females.
6. Early-life food shortage affected very few traits, except for a weak interaction with temperature that affected total fecundity. At 28°C, females that experienced early-life food restriction produced fewer offspring than females with continual food supply. No such diet effect occurred at 24°C.
7. Our results suggest that tropical fish may be severely impacted by increased temperatures (i.e., decreased reproduction with increased morality), but are likely to be resilient to brief periods of food limitations during early development.
8. Interestingly, early-life food shortage caused a reduction in total offspring number but only at 28°C, suggesting that global prey decline might exacerbate the negative effects of a warming climate on stock recruitment of tropical fish.

## INTRODUCTION

Mean global temperature, recorded since 1850, reached its highest point in 2023 (C3S, 2024). This rise in temperature has accelerated development and led to the earlier onset of reproduction in ectotherms (Wang et al., 2020). A benefit of this faster life history is that individuals can allocate more into reproduction when young (Niu et al., 2023; Roff, 1992; Wootton et al., 2022). However, a potential cost of this faster life history is that rapid growth and earlier breeding often divert energy away from self-maintenance, accelerating senescence (e.g., telomere shortening) and/or reducing lifespan (Monaghan & Ozanne, 2018; Munch & Salinas, 2009). There is a trade-off between life history components because an individual with a limited energy budget cannot invest maximally into all traits. The total energy budget depends on resource acquisition and assimilation (van Noordwijk & de Jong, 1986). A change in food availability can alter observable trade-offs due to shifts in optimal allocation because of changes in the costs/benefits of specific traits or processes (Descamps et al., 2016). For example, when food is scarce, animals often forgo costly reproduction to improve their survival (Ruf et al., 2006); conversely, in animals with an abundant food supply, reproduction sometimes has no detectable effects on survival (Ricklefs & Cadena, 2007). Climate warming is reducing prey availability in many ecosystems, especially those in already-warm regions (Beaugrand et al., 2003; Boyce et al., 2010; Suikkanen et al., 2013). Fewer available prey raises questions about the combined effects of higher temperatures and lower food availability on the life histories of tropical ecotherms.

Food availability as a juvenile can have immediate effects on physiological and morphological traits during development. If these effects persist into adulthood, they could play a crucial role in determining an individual’s fitness (Klepsatel et al., 2020). For example, insufficient food during development can slow growth and reduce final adult size, leading to increased predation risk and higher mortality (Le Pape & Bonhommeau, 2015). Food is rarely abundant in nature, however, and periods of starvation are common. Many species have therefore evolved to accommodate the likelihood of a lack of food during development and show plasticity to optimise their life-history to the prevailing circumstances. For example, when food is scarce, animals often delay maturation until they reach the normal adult size (Danielsen et al., 2013); or they compensate for slower development by rapidly accelerating their growth if food again becomes available (Hector et al., 2012; Zhu et al., 2016). The obvious cost of a prolonged juvenile stage is to delay the onset of breeding, but the costs of accelerated growth is less apparent and might only be detectable later in life (Metcalfe & Monaghan, 2001; Monaghan & Ozanne, 2018).

It is a challenge to determine if there is a long-term cost of low food availability during development when it covaries with temperature. For example, low food availability might improve mitochondrial performance and trigger repair pathways (Mattson et al., 2014), potentially improving future tolerance to heat stress. Conversely, some bioenergetic models predict that lower food intake reduces upper thermal limits for ectotherms that are in already-warm environments due to metabolic meltdown (Huey & Kingsolver, 2019). For instance, more than ten ectothermic species prefer to lower their body temperature when they are food restricted (Angilletta, 2009). Higher temperatures tend to elevate metabolism that increases energy loss (Schulte, 2015), or to limit the time spent foraging that then reduces energy intake (Sinervo et al., 2010). These metabolic and behavioural changes, when combined with low food availability, should increase the likelihood of starvation (Kawaguchi et al., 2007). However, ectotherms that acclimate to warmer environments as juveniles might mitigate the costs of an elevated metabolism (Sandblom et al., 2014), or they might lower their metabolic rate when less food is available (Auer et al., 2015). It is still unclear whether or not high temperatures alter the usual life history response of juveniles to lower than normal food availability (e.g., delayed maturation); and, if so, how this affects adult reproductive success and survival.

It is important to investigate how females respond to changes in temperature and food availability, because fewer and/or less fecund females slow population growth (Gownaris & Boersma, 2019). In fish, females with greater adult growth are usually favored by selection because larger females are more fecund (Barneche et al., 2018). More generally, greater investment in somatic maintenance by females than males is expected to make them more resistance to environmental stress (Mauvais-Jarvis, 2024). Intriguingly, several studies have shown that eggs are less sensitive than sperm to heat stress (Iossa, 2019). However, females often require more resources than males to reach and maintain a larger adult size, which could make them more sensitive to low food availability. This conjecture is supported by a recent meta-analysis indicating that females are more sensitive than males to food stress during development (Teder & Kaasik, 2023). Given these contrasting results it is apparent that we need more data on the life-history responses of females to lower food abundance under climate warming. This is particularly relevant for tropical species because rising temperatures should impose greater developmental costs on aquatic ectotherms at lower latitudes (Marshall et al., 2020): historic ambient temperatures in the tropics are closer to those for optimal development than those in temperate areas.

Here, we manipulated both temperature and early-life food availability in a tropical fish – the Trinidadian guppy (*Poecilia reticulata*). Guppies with a higher metabolic rate tend to grow faster, reproduce earlier, and die younger (i.e., have a ‘faster’ pace of life; Auer et al., 2018).

Given that higher temperatures increase metabolic rates in ectotherms, a faster pace of life is a likely life history response by guppies that experience higher temperatures. The water temperature in Trinidad ranges from 20 to 28 °C (Alkins-Koo, 2000). We placed newborn guppies in either control or high temperature rooms (water temperatures of 24.4 ± 0.6 °C and 27.8 ± 0.4 °C) throughout their life, and provided them with either a control or restricted diet between 3 and 17 days of age to mimic a temporary food shortage. A return to a control diet after food restriction causes accelerated juvenile growth in guppies (Auer et al., 2010; Moura-Campos et al., 2024). Once females matured, we paired them with stock males and monitored their adult growth, mortality, and reproductive output for 16 weeks. We then measured a suite of traits related to somatic maintenance and residual reproductive potential. We predicted that any warming-driven shifts in life histories would be more advantageous for females that always had sufficient food because of the greater energy demands of accelerating growth at higher temperatures. We therefore predicted that low food availability early in life would be more detrimental at 28°C than 24°C.

## METHODS

### Origin and maintenance of fish

Stock guppies originating from Alligator creek near Townsville (Australia) were obtained from two separate stock populations (Kranz et al., 2018; Lindholm et al., 2014). This mixed stock population has been kept at the Australian National University since 2019. Stock fish were housed in single-sex 90L aquaria (∼50 fish/tank) at 26 ± 1 °C room temperature under a 14:10 h photoperiod. We fed stock fish twice daily, once with *Artemia* nauplii and once with commercial fish flake.

### Experimental overview

To generate experimental fish, we randomly paired a virgin stock female with a stock male in a 4L tank (*n* = 29 pairs). After two weeks, we removed the males and monitored the females daily for newborn fry. Parents were kept at 24°C and fed *ad libitum* with *Artemia* nauplii twice daily. Newborn fry from each family were alternately assigned to one of four treatments (maximum 10 fry per family/treatment): (a) control temperature-control diet; (b) control temperature-restricted diet; (c) high temperature-control diet; and (d) high temperature-restricted diet (*n* = 110 per treatment). At birth (Day 0), newborn fish were individually transferred to a 1L tank and placed in either high (30 ± 1°C) or control (26 ± 1°C) temperature rooms, resulting in 27.8 ± 0.4 °C and 24.4 ± 0.6 °C water temperature, respectively. Hereafter we refer to the high temperature as 28 °C and low temperatures as 24°C. The assigned temperatures were maintained throughout their life.

We wanted to manipulate food availability shortly after birth when fry growth rates are maximal (Mousavi-Sabet et al., 2014). Some offspring still have remnant egg yolk at birth (Shahjahan et al., 2013), which provides nutrients, so we started the diet treatment three days after birth. On Days 0 to 2, juveniles were fed twice daily with *Artemia* nauplii *ad libitum*. From Days 3 to 16, fish on the control diet were fed *ad libitum* with *Artemia* nauplii twice daily, whereas fish on the restricted diet were fed with 3 mg of wet mass *Artemia* nauplii once every second day. This diet resulted in almost no juvenile growth in another poeciliid species (Vega-Trejo et al., 2016) and in our study (Moura-Campos et al., 2024) without elevating mortality. On Day 17, all fish were returned to the control diet (i.e., *Artemia* nauplii *ad libitum* twice daily). From Day 31, we inspected the fish twice a week to check for maturation (a visible gravid spot). We obtained 217 adult females (*n* = 58 control temperature-control diet; *n* = 52 control temperature-restricted diet; *n* = 55 high temperature-control diet; *n* = 52 high temperature-restricted diet).

Two weeks after maturation, each female was paired with a stock male for 24 hours, then returned to her individual tank to give birth. Males constantly attempt to mate with females and sperm transfer is likely to occur whenever a male is present for 24 hours. This breeding procedure continued each week for 16 weeks – a commonly used period to assess adult performance in guppy studies (Auer, 2010; Auer et al., 2010). Stock males were paired a maximum of once a week and were otherwise kept in male-only stock tanks at 24°C.

### Adult growth

We measured standard length (SL: the snout tip to the end of the vertebral column) of each female at the first mating event (i.e., two weeks after maturation; Week 2), and then every four weeks until the end of the mating period (Week 6, 10, 14 and 18). Females were anaesthetized with *Aqui-S* (0.0075% v/v) for 30 seconds, then placed on their side on a glass slide alongside a scale, and photographed for later measurement of SL using *ImageJ*.

### Reproductive output

Female tanks were monitored daily and newborn offspring were collected. We recorded how many broods each female produced, the date (to estimate gestation duration), the number and size of offspring, and the SL of the female when she gave birth. Female SL was measured using the method outlined in *Adult growth*. To measure offspring size, we placed each newborn fry (*n* = 5358) in a shallow container of tank water and photographed it from above to later measure its length using *ImageJ*.

After 16-weeks, females were euthanized then dissected under a microscope. We counted the number of embryos and unfertilized eggs to assess female residual reproductive potential.

### Mortality and somatic maintenance

We inspected the females every day for 16 weeks to determine the mortality rate. We used the survivors to measure relative telomere length (RTL), immune response, and gut length.

### Relative telomere length (RTL)

Telomere shortening contributes to ageing and is often associated with faster growth, greater reproductive effort, and being in a harsher environment (Chatelain et al., 2020; Monaghan & Ozanne, 2018). We randomly sampled 30 females per treatment (all 18 weeks post-maturation) to measure RTL. A section of tail muscle (20 mg) located between the dorsal and caudal fins was collected and stored at -80 °C. We extracted genomic DNA using DNeasy Blood and Tissue Kit (QIAGEN), and quantified its concentration using Qubit™ dsDNA BR Assay Kits (Thermo Fisher Scientific). DNA purification was checked using a CLARIOstar^®^ Plus microplate reader (BMG Labtech). Real-time quantitative PCR (QuantStudio^®^3 qPCR system) was used to determine RTL by measuring how the telomere DNA sample differed from a non-variable reference DNA sample (Melanocortin 1 receptor; MCR1). We followed the qPCR protocol and primers in two recent studies (Monteforte et al., 2020; Morbiato et al., 2023). Each sample was run in triplicate; and each plate had two inter-plate calibrator samples, and a negative control.

Standard curves were generated by fourfold serial dilutions, which were run in triplicate. The crossing threshold (Ct) values of samples were highly repeatable (*r* ± SE, telomere: 0.989 ± 0.002; MC1R: 0.994 ± 0.001; both *P* = 0.01, *n* = 150). RTL was calculated using the mean Ct values per fish and the equation in (Monteforte et al., 2020).

### Immune response

We used a phytohemagglutinin injection assay to measure cell-mediated immunity (Iglesias-Carrasco et al., 2019; Petitjean et al., 2021). We anaesthetized females and measured their body thickness at the posterior end of the dorsal fin with a pressure-sensitive spessimeter (Mitutoyo 547-301; accuracy: 0.01 mm; five measurements/fish). At the same location, we then injected 0.01 mg of phytohemagglutinin dissolved in 0.01 mL of PBS solution into the side of the fish.

After 24 hours, the body thickness was again measured five times. We calculated the difference in the mean pre-injection and post-injection measures (repeatability: *r* ± SE = 0.983 ± 0.001, *P* = 0.01, *n* = 352) (i.e., swelling) as an index of immunity.

### Gut length

A longer gut can increase nutrient absorption and prolong food retention time (Cant et al., 1996), but there might be greater maintenance costs (Hammond & Diamond, 1992), diverting energy from other traits (Kotrschal et al., 2013). Three days after the immunity test, females were fasted for 24 hours to empty their gut (Kotrschal et al., 2013). We then dissected the females to extract and photograph the gut (the entire digestive tract from mouth to anus), the length of which was measured using *ImageJ*.

## Statistical analyses

Each trait was analyzed separately using generalized linear mixed models (GLMMs; *glmmTMB* package) with the appropriate error distribution (Table S1). Dispersion tests (*DHARMa* package) were performed to confirm that the data variance met the model assumptions. In all analyses, temperature (24°C, 28°C) and early-life food availability (control, restricted) were considered as fixed factors, with their two-way interaction included in initial models. Family identity was considered as a random factor to control for measurements of multiple females from the same family. For models accounting for zero-inflation (Table S1), all predictors in the conditional part of the models were included in the zero-inflation part, except for immune response, where we used an intercept-only model for the zero-inflation part due to convergence issues.

We included standardized SL (mean = 0; SD = 1) as a covariate to estimate the gut length relative to body size. We log-transformed both gut length and SL to account for their positive allometric relationship (Figure S3). We included pre-injection measurement of body thickness (standardized) as a covariate for immune response because females with a thicker body might experience more swelling after the PHA injection.

Wald chi-square tests were conducted to calculate *P* values using the *car* package (*Anova* function). Type III sums of squares were used for models with the interaction term, and type II sums of squares for models without the interaction. If there was a significant interaction, we ran Tukey’s post-hoc pairwise tests (*emmeans* package) to compare different temperature-food groups. Non-significant interactions were removed to interpret main effects (Engqvist, 2005). Statistical outputs of initial and final models are in the supplementary material. Results are presented as mean ± *SE*. The significance level is *α* = 0.05 (two tailed). Analyses were run in R studio v1.3.1093 with R v4.0.5. Any deviations from this approach are specified below.

### Adult growth

We considered SL as the response variable and included adult age (in weeks) and age^2^ to the model as covariates to account for non-linear growth. Female identity was included as a random factor to control for repeated measurements of each female. The initial model included a three-way interaction among temperature, food availability and age. Due to a significant three-way interaction, we ran separate GLMMs for each age to examine the temperature×food interactive effect on SL.

### Reproductive output across broods

To test if females adjusted their reproductive investment (gestation time, offspring number and size) in relation to how many times they had given birth, we included brood order as a fixed factor, and examined its three-way interaction with temperature and food availability. Very few females had more than four broods (5 of 217; all from 28°C), so we only considered the first four broods. Female identity was included as a random effect to account for repeated measurements of each female. Non-significant three-way interactions were excluded to explore two-way interactions (temperature×food, temperature×brood, food×brood).

For offspring size, we assessed both the average size and the coefficient of variation (CV) within a brood. The CV reflects developmental variation because of environmental or maternal effects (Morrongiello et al., 2012; Rowiński & Rogell, 2017). For the average offspring size, the SL of each offspring was the response variable, with brood identity included as a random factor to correct for multiple offspring per brood. For the within-brood variation in offspring size, we estimated Bao’s CV using log-transformed data (Zheng et al., 2023) to account for brood differences in sample sizes and data distributions (Yang et al., 2020). We could use only broods with four or more offspring (*n* = 457 of 526 broods) to calculate Bao’s CV.

### Mortality

We used the stratified log-rank test (*survival* package) to compare survival curves among the four temperature-food treatments. With a significant treatment difference, we compared the survival curves between 24 and 28°C, as well as between the control and restricted diet groups at each temperature. We considered family identity as a stratum and treated individuals that were alive at the end of the mating period as right-censored data.

### Effect of female size on trait expression

A significant treatment effect on some traits might be attributed to the treatment effect on body size (Nakagawa et al., 2017). The intermediate variable of size might then explain differences in other traits (e.g., larger females produce more eggs). To test for size-mediated treatment effects, we ran a second set of analyses that included absolute SL (standardized) as a covariate. For the reproductive traits, we used the SL when a female gave birth; for the other traits, we used the SL at the end of the mating period. We excluded total brood number or total offspring number from these analyses because they represent cumulative values over time without corresponding body size. If significant main effects became non-significant after correcting for body size, it suggests that the effects of temperature and/or food availability mainly resulted from how they affected female size.

## RESULTS

### Adult growth

There was a significant three-way interaction among temperature, food availability and adult age on female size (*χ*^2^1 = 5.249; *P* = 0.022). Temperature affected female size, but this effect depended on adult age (Figure 1*A*). During early adulthood, females at 28°C were significantly larger than those at 24°C (Weeks 2 and 6: both *χ*^2^1 > 9.952; *P* < 0.003). However, adult growth was slower at 28°C, with females at both temperatures reaching the same size by Week 10 (*χ*^2^1 = 0.246; *P*= 0.620). By Week 14, females were significantly smaller at 28°C than 24°C (weeks 14 and 18: both *χ*^2^1 > 14.374; *P* < 0.001). Early-life food availability did not affect adult size, either on its own, or in interaction with temperature (all ages: *χ*^2^1 < 3.168; *P* > 0.074) (Table S2).

**Figure 1.**
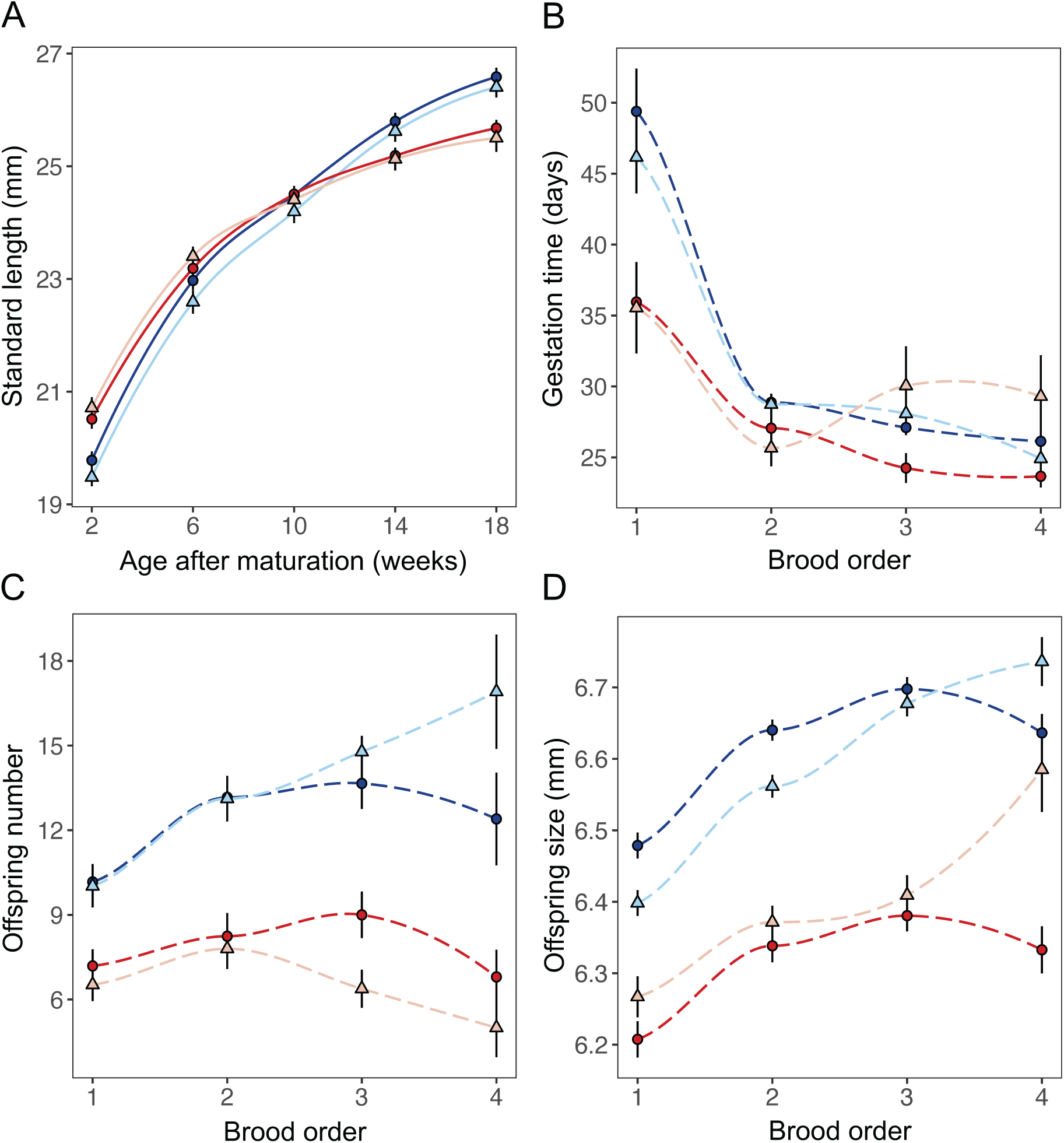
Effects of temperature and early-life food availability on female size over time and reproductive investment across broods. Colors indicate temperature (24°C = blue; 28°C = red). Darkness-shapes indicate early-life food availability (control diet = dark circle; restricted diet = light triangle). Line bars show mean ± *SE*.

### Reproductive output

Temperature and brood order independently affected offspring size, but they interacted to affect gestation time, offspring number per brood, and within-brood variation in offspring size (Table 1). Early-life food availability did not affect any of the above reproductive traits, either independently or in interaction with temperature or brood order (Table 1). The only exception was that early-life food availability affected overall fecundity (Table 2).

**Table 1.**
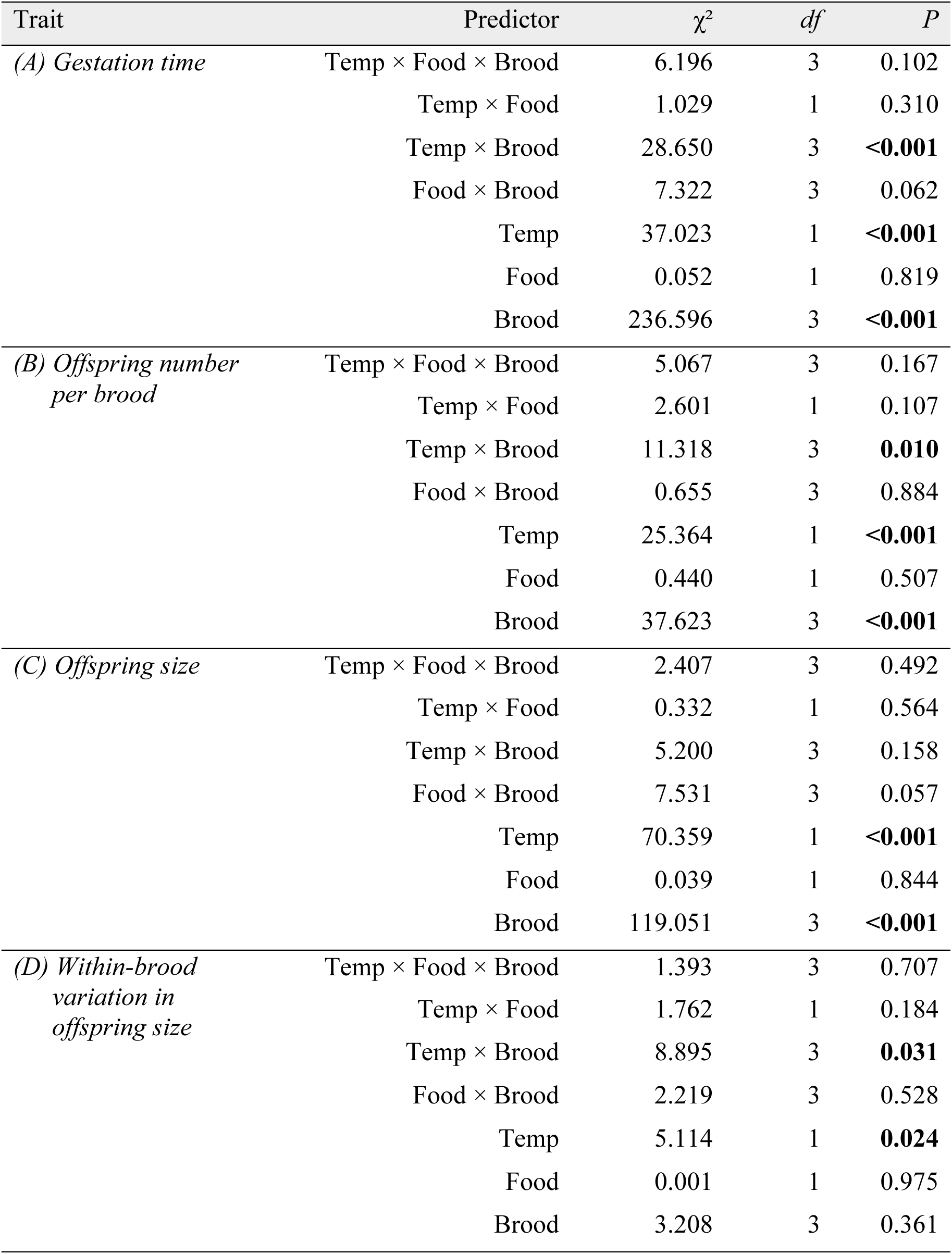
Effects of temperature, early-life food availability, and brood order on female reproductive investment in successive broods. *P*-values are from models excluding non-significant higher order interactions. Outputs of the full models are in the supplementary material (Table S3).

| Trait | Predictor | $\chi^2$ | *df* | *P* |
| --- | --- | --- | --- | --- |
| *(A) Gestation time* | Temp × Food × Brood | 6.196 | 3 | 0.102 |
|  | Temp × Food | 1.029 | 1 | 0.310 |
|  | Temp × Brood | 28.650 | 3 | **<0.001** |
|  | Food × Brood | 7.322 | 3 | 0.062 |
|  | Temp | 37.023 | 1 | **<0.001** |
|  | Food | 0.052 | 1 | 0.819 |
|  | Brood | 236.596 | 3 | **<0.001** |
| *(B) Offspring number per brood* | Temp × Food × Brood | 5.067 | 3 | 0.167 |
|  | Temp × Food | 2.601 | 1 | 0.107 |
|  | Temp × Brood | 11.318 | 3 | **0.010** |
|  | Food × Brood | 0.655 | 3 | 0.884 |
|  | Temp | 25.364 | 1 | **<0.001** |
|  | Food | 0.440 | 1 | 0.507 |
|  | Brood | 37.623 | 3 | **<0.001** |
| *(C) Offspring size* | Temp × Food × Brood | 2.407 | 3 | 0.492 |
|  | Temp × Food | 0.332 | 1 | 0.564 |
|  | Temp × Brood | 5.200 | 3 | 0.158 |
|  | Food × Brood | 7.531 | 3 | 0.057 |
|  | Temp | 70.359 | 1 | **<0.001** |
|  | Food | 0.039 | 1 | 0.844 |
|  | Brood | 119.051 | 3 | **<0.001** |
| *(D) Within-brood variation in offspring size* | Temp × Food × Brood | 1.393 | 3 | 0.707 |
|  | Temp × Food | 1.762 | 1 | 0.184 |
|  | Temp × Brood | 8.895 | 3 | **0.031** |
|  | Food × Brood | 2.219 | 3 | 0.528 |
|  | Temp | 5.114 | 1 | **0.024** |
|  | Food | 0.001 | 1 | 0.975 |
|  | Brood | 3.208 | 3 | 0.361 |

**Table 2.**
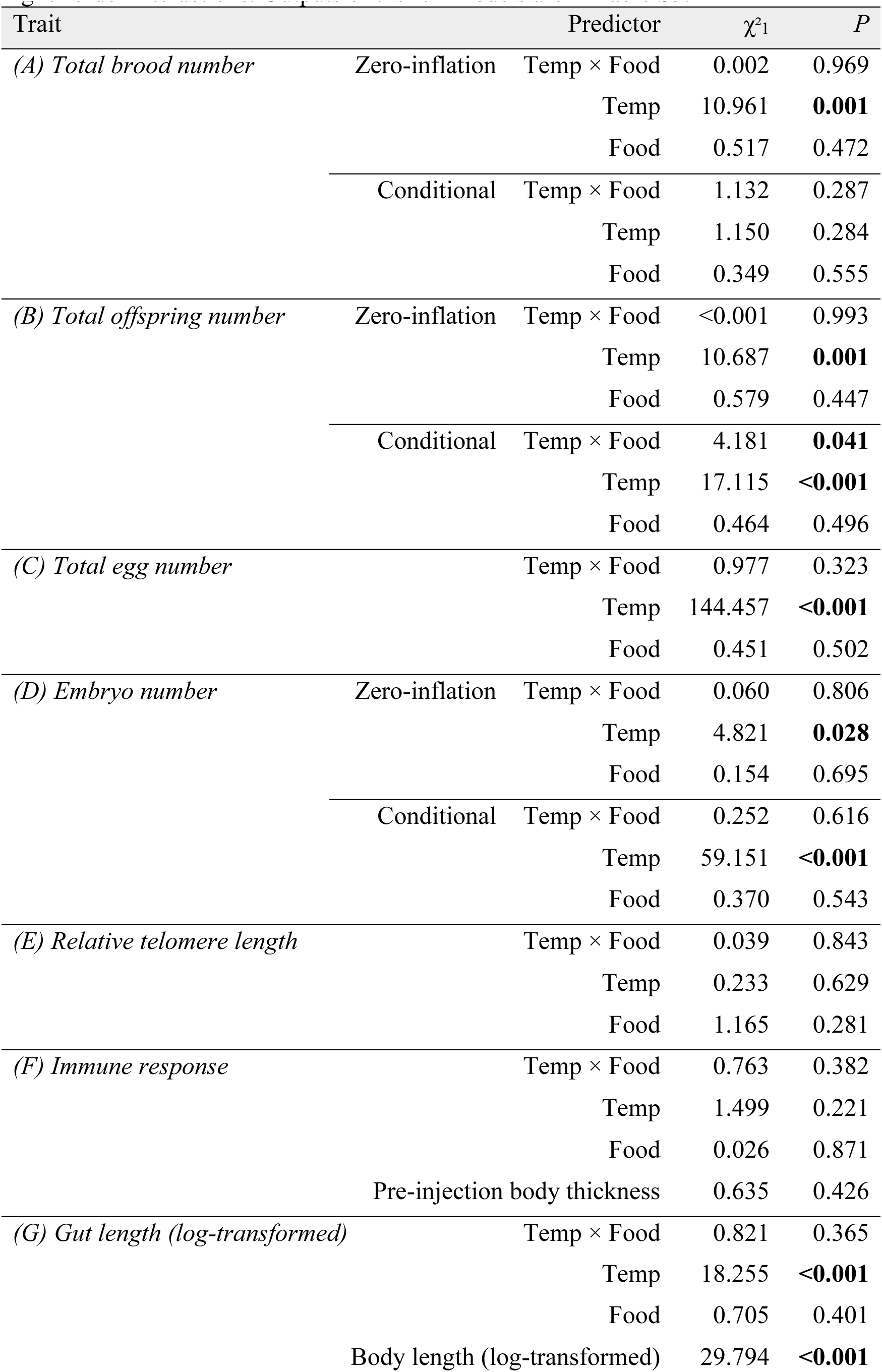
Effects of temperature and early-life food availability on female overall fecundity and somatic investment. *P*-values are from models excluding non-significant higher order interactions. Outputs of the full models are in Table S3.

| Trait | | Predictor | $\chi^2_1$ | *P* |
| --- | --- | --- | --- | --- |
| *(A) Total brood number* | Zero-inflation | Temp × Food | 0.002 | 0.969 |
|  |  | Temp | 10.961 | **0.001** |
|  |  | Food | 0.517 | 0.472 |
| | Conditional | Temp × Food | 1.132 | 0.287 |
|  |  | Temp | 1.150 | 0.284 |
|  |  | Food | 0.349 | 0.555 |
| *(B) Total offspring number* | Zero-inflation | Temp × Food | <0.001 | 0.993 |
|  |  | Temp | 10.687 | **0.001** |
|  |  | Food | 0.579 | 0.447 |
| | Conditional | Temp × Food | 4.181 | **0.041** |
|  |  | Temp | 17.115 | **<0.001** |
|  |  | Food | 0.464 | 0.496 |
| *(C) Total egg number* | | Temp × Food | 0.977 | 0.323 |
|  |  | Temp | 144.457 | **<0.001** |
|  |  | Food | 0.451 | 0.502 |
| *(D) Embryo number* | Zero-inflation | Temp × Food | 0.060 | 0.806 |
|  |  | Temp | 4.821 | **0.028** |
|  |  | Food | 0.154 | 0.695 |
| | Conditional | Temp × Food | 0.252 | 0.616 |
|  |  | Temp | 59.151 | **<0.001** |
|  |  | Food | 0.370 | 0.543 |
| *(E) Relative telomere length* | | Temp × Food | 0.039 | 0.843 |
|  |  | Temp | 0.233 | 0.629 |
|  |  | Food | 1.165 | 0.281 |
| *(F) Immune response* | | Temp × Food | 0.763 | 0.382 |
|  |  | Temp | 1.499 | 0.221 |
|  |  | Food | 0.026 | 0.871 |
|  |  | Pre-injection body thickness | 0.635 | 0.426 |
| *(G) Gut length (log-transformed)* | | Temp × Food | 0.821 | 0.365 |
|  |  | Temp | 18.255 | **<0.001** |
|  |  | Food | 0.705 | 0.401 |
|  |  | Body length (log-transformed) | 29.794 | **<0.001** |

### Gestation time

Females at both temperatures took significantly longer to give birth to their first brood than their subsequent broods (Figure 1*B*; Table S3*A*). Females at 28°C had their first brood sooner than females at 24°C (Tukey’s test: *P* <0.001), but the gestation time for subsequent broods did not differ between the two temperatures (all *P* > 0.246; Figure 1*B*).

When female size was controlled for (*χ*^2^1 = 208.679; *P* < 0.001), females at 28°C still had their first brood significantly sooner than females at 24°C (Tukey’s test: *P* < 0.001), but the gestation time for third and fourth broods was significantly longer at 28°C than 24°C (both *P* < 0.027).

### Offspring number per brood

Across all four broods, females at 28°C produced significantly fewer offspring per brood than females at 24°C (Tukey’s test: all *P* <0.001; Figure 1*C*). At 24°C, females produced significantly fewer offspring in the first brood than in subsequent broods (all *P* <0.002), which each contained a similar number of offspring (all *P* > 0.564). At 28°C, however, the number of offspring was similar across all four broods (all *P* > 0.073).

When female size was controlled for (*χ*^2^1 = 25.578; *P* < 0.001), there was no longer an interaction between temperature and brood order affecting the number of offspring (*χ*^2^3 = 6.788; *P* = 0.079). However, the number of offspring remained significantly lower at 28°C than at 24°C (*χ*^2^1 = 83.762; *P* < 0.001).

### Offspring size

Offspring of females that developed at 28°C were significantly smaller than those of females that developed at 24°C, irrespective of brood order (Table 1*C*; Figure 1*D*). In addition, at both temperatures, first-brood offspring were significantly smaller than those from subsequent broods (Tukey’s test: all *P* < 0.001). In contrast, an interaction between temperature and brood order affected within-brood variation in offspring size (Table 1*D*). Offspring size in the first brood was significantly more variable at 28°C than 24°C (Tukey’s test: *P* = 0.024), with no temperature effect on variability in the next three broods (all *P* > 0.093). Controlling for female size did not alter the above findings (Table S3*C,D*).

### Overall fecundity

Females at 28°C were less likely to give birth than those at 24°C, with no effect of early-life food availability (Table 2*A*). For females that gave birth, the number of broods they produced was unaffected by temperature or food availability (Table 2*A*; Figure 2*A*). However, the total number of offspring produced was significantly lower at 28°C than at 24°C (Figure 2*B*), and lower food availability early in life amplified this effect (interaction: *χ*^2^1 = 4.181; *P* = 0.041) (Table 2*B*), resulting in a further reduction in total offspring number at 28°C (Tukey’s test: *P* = 0.044), but not at 24°C (*P* = 0.497).

**Figure 2.**
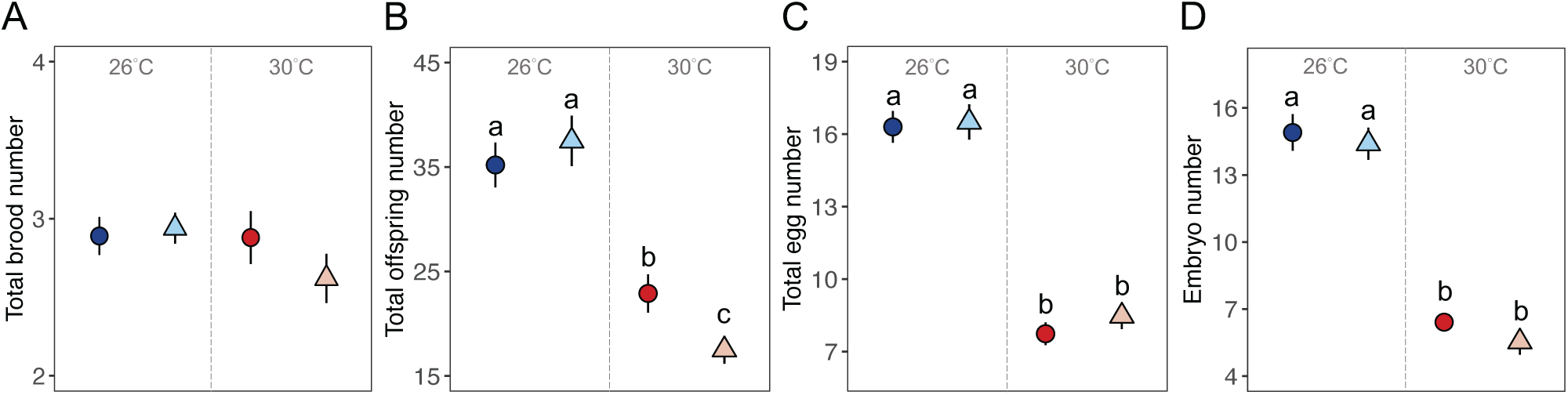
**Effects of temperature and early-life food availability on overall fecundity**. (A) total brood number, (B) total offspring number, (C) total egg number, (D) embryo number. Total brood number and total offspring number were values accumulated over the entire mating period, while total egg number and embryo number were measured at the end of the period. Figures show mean ± *SE* and only present the conditional part of the results (i.e., exclude zero values) (see Table 2 for details). Different temperatures are presented using color (24°C = blue; 28°C = red), and different food availability are shown using darkness-shape (control diet = dark circle; restricted diet = light triangle). Letters indicate significant differences between treatments using Tukey’s tests.

At the end of the mating period, females at 28°C held fewer eggs and fewer embryos than those at 24°C (Table 2*C,D*), with no effects of early-life food availability (Figure 2*C,D*). When female size was controlled for, the effects of temperature on the number of eggs or embryos remained significant (Table S4*C,D*).

### Mortality and somatic investment

Survival curves differed significantly among the four treatments (Figure 3). Temperature significantly affected adult female mortality (*χ*^2^1 = 44.400; *P* < 0.001): at 28°C in total 43 of 107 females died, compared to only 1 of 109 females at 24°C. At both temperatures, however, early-life food availability did not affect morality (both *χ*^2^1 < 0.700; *P* > 0.400).

**Figure 3.**
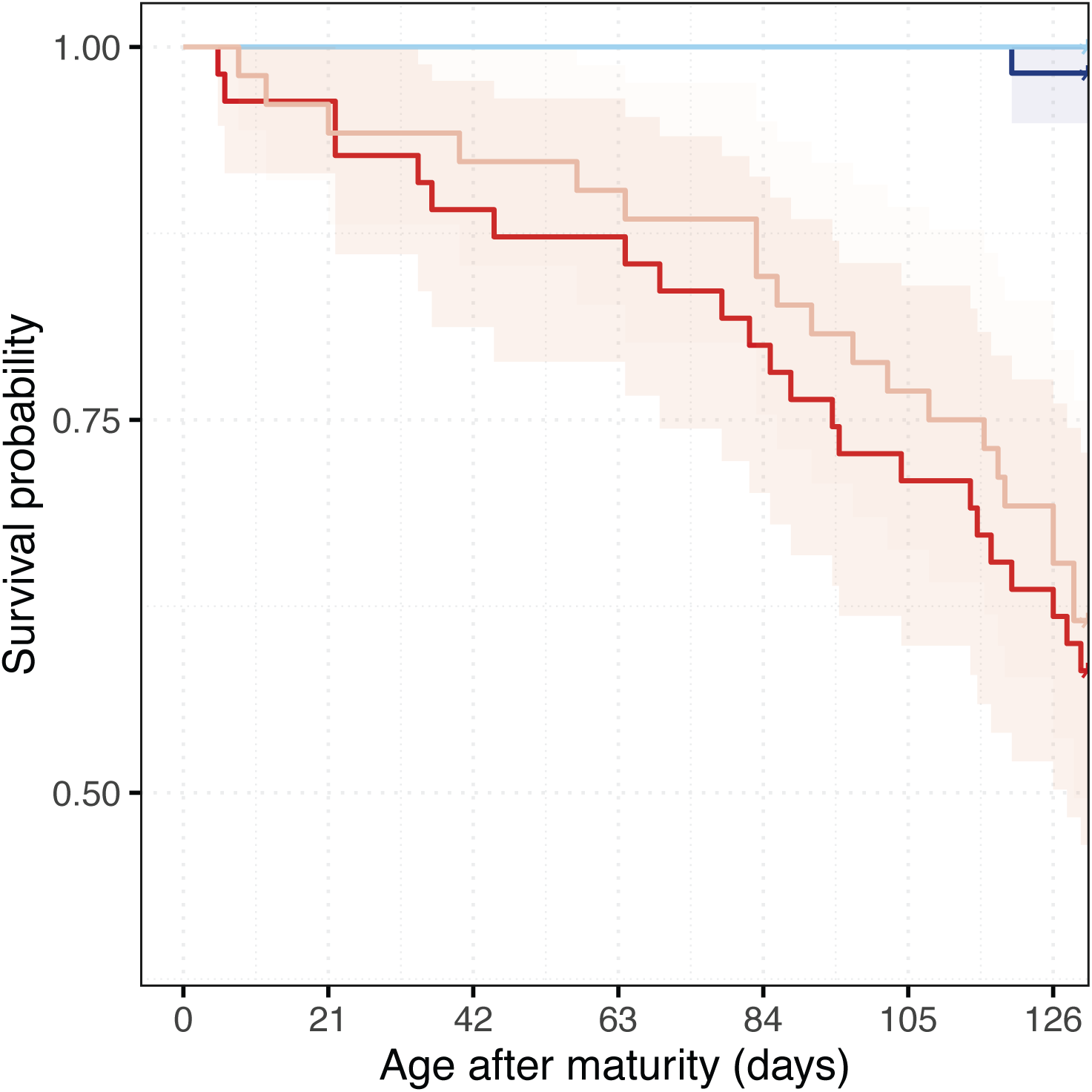
**Effects of temperature and early-life food availability on mortality**. Treatment is presented using colors: 24°C-control diet (dark blue), 24°C-restricted diet (light blue), 28°C-control diet (dark red), 28°C -restricted diet (light red) along with 95% confidence intervals.

For females that survived until the end of the experiment, their RTL and immune response was unaffected by temperature, early-life food availability, or their interaction (Table 2; Figure 4); these results remained unchanged after controlling for female size (Table S5). RTL was unrelated to chronological age (i.e., birth to age of testing: *χ*^2^1 = 2.045; *P* = 0.153). Interestingly, females at 28°C had a significantly longer gut (corrected for body size) than those at 24°C (Table 2G), with no effect of early-life food availability (Figure 4C).

**Figure 4.**
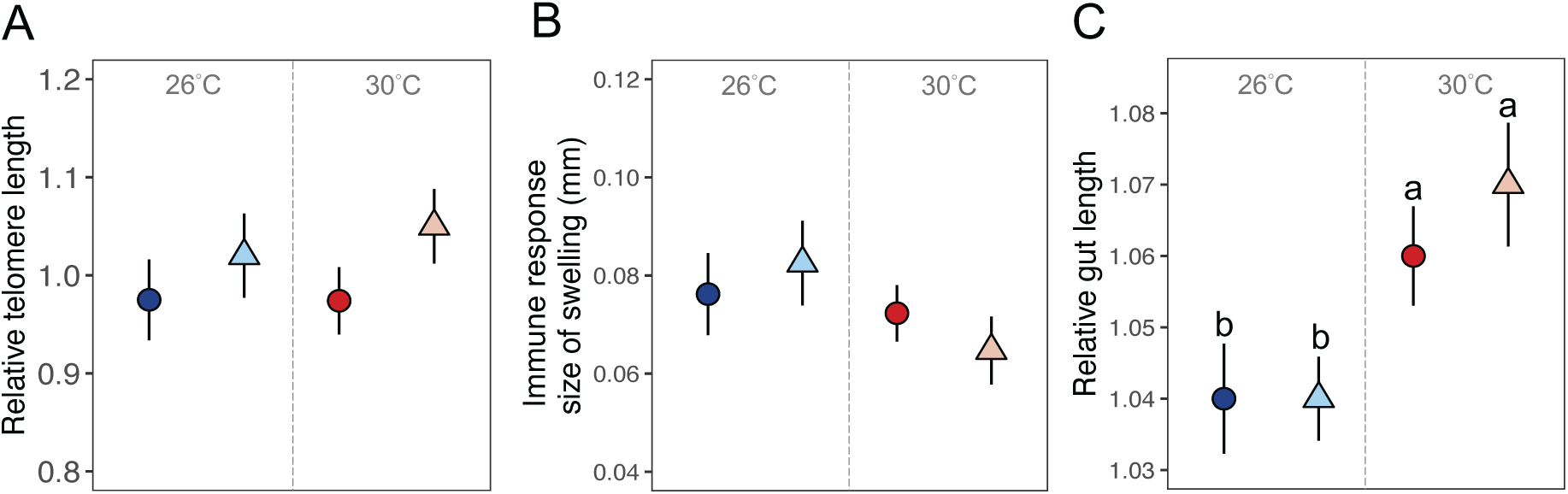
Effects of temperature and early-life food availability on somatic investment. (A) Relative telomere length, (B) immunity, (C) relative gut length. Colors indicate temperature (24°C = blue; 28°C = red). Darkness-shapes indicate food availability (control diet = dark circle; restricted diet = light triangle). Figures show mean ± *SE.* Letters indicate significant differences using Tukey’s tests.

## DISCUSSION

Natural selection generates optimal life histories for past environments (Ghalambor et al., 2007). Novel environments can therefore lead to maladaptive responses. Rapid environmental change imposes novel challenges, such as the co-occurrence of rising temperatures and periods of low food availability that is now affecting tropical ectotherms (Huey & Kingsolver, 2019). Ectotherms often show a phenotypically plastic response to ‘live fast, die young’ in warmer environments (Brans & De Meester, 2018; Mundinger et al., 2022). But is this response possible when food is limited during early development? And is this ‘fast’ life history adaptive (Bestion et al., 2015; Wang et al., 2020)? Female guppies that experience early-life food restriction adjust their life history by accelerating their juvenile growth and changing the time and size at maturation when they return to a normal diet (Auer, 2010). Recent work shows that these plastic developmental responses are temperature-dependent (Moura-Campos et al., 2024), but it is unclear how these life history shifts affect net reproductive output, and whether this is also temperature-dependent.

In our study, temperature strongly affected female life histories. Irrespective of their early diet, female guppies at the higher temperature grew more slowly as adults, reproduced less often and died younger. In contrast, there was no effect of food availability on any of the measured traits. The only exception is that early-life food restriction reduced the total number of offspring at 28°C but not 24°C. Even here, however, the strength of the effect of food availability was very small compared to the effect of temperature on a range of traits (see Tables 1 and 2). We conclude that: (1) female guppies are generally resilient to a period of food restriction early in life and (2) temperature has a much bigger influence on their life histories.

### A higher temperature led to a fast life history with lower reproductive output

Our results generally support the pattern of a shift to ‘live fast, die young’ at higher temperatures: female guppies reproduced earlier and died younger at 28°C than 24°C. However, a higher temperature did not cause greater reproductive investment by guppies in early than late adulthood (Niu et al., 2023; Roff, 1992). Females consistently produced broods with fewer and smaller offspring at 28°C than 24°C. As a result, net reproductive success (total offspring number) was lower at 28°C. The adaptive value of an earlier onset of reproduction at the higher temperature is therefore unclear. A trend for females to produce smaller offspring in hotter environments occurs in many taxa (Bownds et al., 2010; Burgess & Marshall, 2011; Fox & Czesak, 2000). This trend could be adaptive because being smaller lowers the total energetic costs of development which are greater due to a higher metabolic rate at hotter temperatures (Gillooly et al., 2001; Riemer et al., 2018). However, smaller offspring also have fewer energy reserves and tend to be more vulnerable to other stressors (e.g., predation, competition) (Aubret, 2012; Wootton & Smith, 2015), which can significantly lower recruitment rates (Houde, 1987).

In our study, the production of smaller offspring at the higher temperature was not associated with having more offspring (i.e., no quality-quantity trade-off; Stearns, 1992). In fact, female guppies also produced fewer offspring at the higher temperature. In contrast, for example, Rutschmann et al. (2016) reported that common lizards produced more offspring at a higher temperature, albeit of the same size as those produced at a lower temperature. The reason for differences between ectotherms in their life history responses to higher temperatures remains unclear.

In guppies, producing fewer offspring at higher temperatures could potentially lower population recruitment rates and increase the risk of local extinctions as the climate continues to warm. This assumes, however, that smaller offspring have the same or lower survival than larger offspring at a higher temperature so that the shift we observed for females to produce smaller offspring is maladaptive. Future studies should clarify how temperature-dependent changes in reproductive investment affect offspring performance, and thereby population dynamics. It is also worth noting that transgenerational effects might influence life histories and buffer any detrimental impact of a higher temperatures that occur in the initially exposed generation, but might be weaker or absent in their grandoffspring (Niu et al., 2023; Salinas & Munch, 2012; Van Doorslaer et al., 2007).

Female fecundity is size-dependent in fish species (Barneche et al., 2018), and higher temperatures typically cause females to mature at a smaller size (temperature-size rule; Forster et al., 2012). Surprisingly, however, we found exceptions to these patterns in guppies. First, females matured as larger adults at 28°C than 24°C, although they then grew more slowly as adults and ended up being smaller after 14 weeks. Second, young females that developed at 28°C were less fecund than young females that developed at 24°C, despite being larger. Controlling for female size (i.e., differences in adult growth) did not alter the findings, indicating that females that developed at 28°C had consistently lower reproductive output than size-matched females that developed at 24°C. This suggests a temperature-induced shift in the size-fecundity relationship. One possible explanation for this shift is that the physiological costs of reproduction (e.g., metabolic loads) increase with temperature (Ginther et al., 2024; Marshall et al., 2020), lowering fecundity given a similar energy budget. Likewise, growth and/or somatic maintenance can be more demanding at higher temperatures (Barneche et al., 2019), reducing the resources that can be allocated to reproduction. Female guppies at 28°C appeared to be thinner and had a less obvious gravid spot than females at 24°C (Figure S4), suggesting that their poorer nutritional state might account for their lower breeding success despite being larger-bodied (Marshall et al., 2018; Rollinson & Rowe, 2016). However, it is noteworthy that we provided adult female guppies with *ad libitum* food (*Artemia* nauplii) twice a day, which should minimize the risk of resource limitation (Descamps et al., 2016). Our results align, however, with experiments on coral reef fish (Donelson et al., 2010) and marine copepods (Lee et al., 2003), where, despite being provided with ample food, females still showed a decrease in reproduction at higher temperatures. A second, related, explanation for the temperature-induced shift in the size-fecundity relationship in guppies is that adult females at 28°C require more food than the amount we provided due to their higher metabolic rate. Following this logic, food availability would need to *increase* with rising temperatures to maintain the same level of fecundity. Unfortunately, this is not the case in many tropical ecosystems where food availability is declining (Lister & Garcia, 2018; Thomas et al., 2017). The role of metabolism in driving temperature-induced shift in the size-fecundity relationship is, however, unclear. For example, Wootton et al. (2022) recently showed that smaller female zebrafish at higher temperatures allocated more into reproduction, without corresponding changes in their metabolic rate. Their results suggest that, instead of physiological constraints, maturing at a small size might be an adaptive response that enables a faster life history at higher temperatures (Wootton et al., 2022; see also White et al., 2022). Without information about rates of food intake and metabolism, however, it is not possible to distinguish between these competing mechanisms to explain why female guppies did not mature at a smaller size at the higher temperature as seen in other species.

Female guppies that developed at the higher temperature reproduced earlier and died sooner. Did they experience accelerated somatic declines because of their faster life history? Intriguingly, there was no evidence that temperature affected telomere length or immune responses. The higher temperature did, however, cause an increase in relative gut length. In ectotherms, energy intake through food consumption can drop dramatically at higher than optimal temperatures (Huey & Kingsolver, 2019). The observed increase in gut length at 28°C might represent an adaptive plastic response that allows females to maintain sufficient digestion and absorption of nutrients (Cant et al., 1996). Consequently, females at 28°C might maintain somatic functions (i.e., telomere length and immunity) because the elongated gut ensures sufficient resource acquisition. These results should be interpreted cautiously, however, because there was far higher female mortality at 28°C. High adult mortality could create a ‘selection bias’ if females with shorter RTL or weaker immunity were more likely to die and absent from our sample of females measured 16 weeks after maturation (Hämäläinen et al., 2014; Salmón et al., 2017). However, even if high mortality at 28°C did selectively remove females, we still observed fewer eggs and embryos (i.e., lower residual reproductive potential) in the surviving females. Hence, any increase in resource acquisition due to an elongated gut did not translate into enhanced late-life fecundity. Of course, it is possible that the reproductive potential of females at 28°C would have been even lower without an elongated gut.

### Food shortages early in life did not have any main effects on female life history

Lower food availability is expected to change the optimal allocation to traits because the net gain per unit invested varies with the absolute level of investment (i.e., costs or benefits are a non-linear function of resources allocated; Descamps et al., 2016). In our study, food restriction during early development did not affect adult size – either at maturity or subsequent adult growth – indicating that female guppies fully compensated for delayed development (Auer et al., 2010). Many studies have, however, noted that even if adult size is unaffected, there could be ‘hidden’ costs of compensatory growth later in life, including telomere shortening (Geiger et al., 2012) and a shorter lifespan (Lee et al., 2013). We did not detect any effect of early-life food availability on these traits. There was, however, a significant effect of early-life food availability on total offspring number, but this was evident only at the higher temperature.

The contribution of early-life food availability to further lowering fecundity at the higher temperature in guppies persisted after controlling for female size. This implies that predictions about the future productivity of fisheries based solely on the effect of higher temperatures might be overestimates because they fail to account for global declines in food availability. Incorrect estimates will occur when stock assessments do not account for changes in size-dependent fecundity and assume a constant relationship across different environments (Maunder & Piner, 2015). Here, we documented a shift in the relationship between female size and total fecundity at 28°C due to a low food availability. At a higher temperature, females that experienced low-food availability early in life had fewer offspring than size-matched females with constant access to food. If the pattern we observed in guppies occurs in other fish species, it will be more challenging to predict fishery dynamics as temperatures rise. For example, the number of larger females might not link to recruitment rates in a fish population (Beldade et al., 2012) as predicted if these females experienced a period of starvation during early development. We note, however, that our evidence is weak as the interaction between early diet and temperatures affecting offspring production by breeding females was only marginally significant (*P* = 0.04), and there was no equivalent interaction affecting whether or not females produced any offspring. To draw more robust conclusions, more data is needed on other fish species.

## CONCLUSION

Our finding that guppies have a fast life history with reduced reproductive output at higher temperatures aligns with findings for marine fishes (Wang et al., 2020). It indicates that fish might not benefit from a faster life-history at higher temperatures. Instead, temperature-dependent changes in life histories might lower population viability (Bestion et al., 2015). In contrast, there was a weak effect of early-life food availability on total fecundity, which was only apparent at a higher temperature. Adult females have indeterminate growth in guppies. It is therefore possible that adult food availability might play an important role because it can offset the immediate effects of a temporary food shortage early in life. This explanation is supported by evidence from other fish that adult diet has a stronger effect than juvenile diet on gametes (Macartney et al., 2019). In sum, when higher temperatures are associated with lower prey abundance, our study suggests that changes in life histories will be most due to temperature rather than early-life food availability.

## SUPPLEMENTARY MATERIAL

Supplementary material has been uploaded as a separate file.

## CODE AND DATA ACCESSIBILITY

Raw data and R code can be downloaded from Dryad https://datadryad.org/stash/share/pYjA-meoq-gX8xpGGTE533ELuOsBU0OSTrZdoSq_RaY

## Acknowledgements

## Acknowledgements

We thank Yusheng Wang, Ruijia Liang, Rozie Higgs and Kate Farkas for the assistance with data collection, and the ANU Animal Services staff for the help with fish maintenance.

## Author contributions

MHJC, DMC and MLH conceived and designed the study. MHJC, CZ and DMC carried out experimental work. MHJC analyzed data, interpreted the results and wrote the draft of the manuscript, with MLH and MDJ providing critical revisions. All authors approved the final manuscript.

## Ethics

The project received approval from the Australian National University’s Animal Ethics Committee (A2021/04).

## REFERENCES

1. Alkins-Koo, M. (2000). Reproductive timing of fishes in a tropical intermittent stream. Environmental Biology of Fishes, 57, 49–66. 10.1023/A:1007566609881

2. Angilletta, M. J. (2009). Thermoregulation. In M. J. Angilletta (Eds.), Thermal Adaptation: A Theoretical and Empiricial Synthesis (pp. 88–125). Oxford University Press.

3. Aubret, F. (2012). Body-size evolution on islands: are adult size variations in tiger snakes a nonadaptive consequence of selection on birth size? American Naturalist, 179, 756–767. 10.1086/665653

4. Auer, S. K. (2010). Phenotypic plasticity in adult life-history strategies compensates for a poor start in life in Trinidadian guppies (*Poecilia reticulata*). American Naturalist, 176, 818–829. 10.1086/657061

5. Auer, S. K., Arendt, J. D., Chandramouli, R., & Reznick, D. N. (2010). Juvenile compensatory growth has negative consequences for reproduction in Trinidadian guppies (*Poecilia reticulata*). Ecology Letters, 13, 998–1007. 10.1111/j.1461-0248.2010.01491.x

6. Auer, S. K., Dick, C. A., Metcalfe, N. B., & Reznick, D. N. (2018). Metabolic rate evolves rapidly and in parallel with the pace of life history. Nature Communications, 9, 14. 10.1038/s41467-017-02514-z

7. Auer, S. K., Salin, K., Rudolf, A. M., Anderson, G. J., & Metcalfe, N. B. (2015). Flexibility in metabolic rate confers a growth advantage under changing food availability. Journal of Animal Ecology, 84, 1405–1411. 10.1111/1365-2656.12384

8. Barneche, D. R., Jahn, M., & Seebacher, F. (2019). Warming increases the cost of growth in a model vertebrate. Functional Ecology, 33, 1256–1266. 10.1111/1365-2435.13348

9. Barneche, D. R., Robertson, D. R., White, C. R., & Marshall, D. J. (2018). Fish reproductive-energy output increases disproportionately with body size. Science, 360, 642–644. 10.1126/science.aao6868

10. Beaugrand, G., Brander, K. M., Lindley, J. A., Souissi, S., & Reid, P. C. (2003). Plankton effect on cod recruitment in the North Sea. Nature, 426, 661–664. 10.1038/nature02164

11. Beldade, R., Holbrook, S. J., Schmitt, R. J., Planes, S., Malone, D., & Bernardi, G. (2012). Larger female fish contribute disproportionately more to self-replenishment. Proceedings of the Royal Society B-Biological Sciences, 279, 2116–2121. 10.1098/rspb.2011.2433

12. Bestion, E., Teyssier, A., Richard, M., Clobert, J., & Cote, J. (2015). Live fast, die young: experimental evidence of population extinction risk due to climate change. PLoS Biology, 13, e1002281. 10.1371/journal.pbio.1002281

13. Bownds, C., Wilson, R., & Marshall, D. J. (2010). Why do colder mothers produce larger eggs? An optimality approach. Journal of Experimental Biology, 213, 3796–3801. 10.1242/jeb.043356

14. Boyce, D. G., Lewis, M. R., & Worm, B. (2010). Global phytoplankton decline over the past century. Nature, 466, 591–596. 10.1038/nature09268

15. Brans, K. I., & De Meester, L. (2018). City life on fast lanes: urbanization induces an evolutionary shift towards a faster lifestyle in the water flea *Daphnia*. Functional Ecology, 32, 2225–2240. 10.1111/1365-2435.13184

16. Burgess, S. C., & Marshall, D. J. (2011). Temperature-induced maternal effects and environmental predictability. Journal of Experimental Biology, 214, 2329–2336. 10.1242/jeb.054718

17. Copernicus Climate Change Service (2024) The 2023 Annual Climate Summary-Global Climate Highlights.

18. Cant, J. P., McBride, B. W., & Croom, W. J. (1996). The regulation of intestinal metabolism and its impact on whole animal energetics. Journal of Animal Science, 74, 2541–2553.

19. Chatelain, M., Drobniak, S. M., & Szulkin, M. (2020). The association between stressors and telomeres in non-human vertebrates: a meta-analysis. Ecology Letters, 23, 381–398. 10.1111/ele.13426

20. Danielsen, E. T., Moeller, M. E., & Rewitz, K. F. (2013). Nutrient signaling and developmental timing of maturation. Current Topics in Developmental Biology, 105, 37–67. 10.1016/B978-0-12-396968-2.00002-6

21. Descamps, S., Gaillard, J. M., Hamel, S., & Yoccoz, N. G. (2016). When relative allocation depends on total resource acquisition: implication for the analysis of trade-offs. Journal of Evolutionary Biology, 29, 1860–1866. 10.1111/jeb.12901

22. Donelson, J. M., Munday, P. L., McCormick, M. I., Pankhurst, N. W., & Pankhurst, P. M. (2010). Effects of elevated water temperature and food availability on the reproductive performance of a coral reef fish. Marine Ecology Progress Series, 401, 233–243. 10.3354/meps08366

23. Engqvist, L. (2005). The mistreatment of covariate interaction terms in linear model analyses of behavioural and evolutionary ecology studies. Animal Behaviour, 70, 967–971. 10.1016/j.anbehav.2005.01.016

24. Forster, J., Hirst, A. G., & Atkinson, D. (2012). Warming-induced reductions in body size are greater in aquatic than terrestrial species. Proceedings of the National Academy of Sciences of the United States of America, 109, 19310–19314. 10.1073/pnas.1210460109

25. Fox, C. W., & Czesak, M. E. (2000). Evolutionary ecology of progeny size in arthropods. Annual Review of Entomology, 45, 341–369. 10.1146/annurev.ento.45.1.341

26. Geiger, S., Le Vaillant, M., Lebard, T., Reichert, S., Stier, A., Le Maho, Y., & Criscuolo, F. (2012). Catching-up but telomere loss: half-opening the black box of growth and ageing trade-off in wild king penguin chicks. Molecular Ecology, 21, 1500–1510. 10.1111/j.1365-294X.2011.05331.x

27. Ghalambor, C. K., McKay, J. K., Carroll, S. P., & Reznick, D. N. (2007). Adaptive versus non-adaptive phenotypic plasticity and the potential for contemporary adaptation in new environments. Functional Ecology, 21, 394–407. 10.1111/j.1365-2435.2007.01283.x

28. Gillooly, J. F., Brown, J. H., West, G. B., Savage, V. M., & Charnov, E. L. (2001). Effects of size and temperature on metabolic rate. Science, 293, 2248–2251. 10.1126/science.1061967

29. Ginther, S. C., Cameron, H., White, C. R., & Marshall, D. J. (2024). Metabolic loads and the costs of metazoan reproduction. Science, 384, 763–767. 10.1126/science.adk6772

30. Gownaris, N. J., & Boersma, P. D. (2019). Sex-biased survival contributes to population decline in a long-lived seabird, the Magellanic Penguin. Ecological Applications, 29, e01826. 10.1002/eap.1826

31. Hämäläinen, A., Dammhahn, M., Aujard, F., Eberle, M., Hardy, I., Kappeler, P. M., Perret, M., Schliehe-Diecks, S., & Kraus, C. (2014). Senescence or selective disappearance? Age trajectories of body mass in wild and captive populations of a small-bodied primate. Proceedings of the Royal Society B-Biological Sciences, 281, 20140830. 10.1098/rspb.2014.0830

32. Hammond, K. A., & Diamond, J. (1992). An experimental test for a ceiling on sustained metabolic rate in lactating mice. Physiological Zoology, 65, 952–977. 10.1086/physzool.65.5.30158552

33. Hector, K. L., Bishop, P. J., & Nakagawa, S. (2012). Consequences of compensatory growth in an amphibian. Journal of Zoology, 286, 93–101. 10.1111/j.1469-7998.2011.00850.x

34. Houde, E. D. (1987). Fish early life dynamics and recruitment variability. American Fisheries Society symposium, 2, 17–29.

35. Huey, R. B., & Kingsolver, J. G. (2019). Climate warming, resource availability, and the metabolic meltdown of ectotherms. American Naturalist, 194, 140–150. 10.1086/705679

36. Iglesias-Carrasco, M., Fox, R. J., Vincent, A., Head, M. L., & Jennions, M. D. (2019). No evidence that male sexual experience increases mating success in a coercive mating system. Animal Behaviour, 150, 201–208. 10.1016/j.anbehav.2019.02.012

37. Iossa, G. (2019). Sex-specific differences in thermal fertility limits. Trends in Ecology & Evolution, 34, 490–492. 10.1016/j.tree.2019.02.016

38. Kawaguchi, Y., Miyasaka, H., Genkai-Kato, M., Taniguchi, Y., & Nakano, S. (2007). Seasonal change in the gastric evacuation rate of rainbow trout feeding on natural prey. Journal of Fish Biology, 71, 1873–1878. 10.1111/j.1095-8649.2007.01647.x

39. Klepsatel, P., Knoblochová, D., Girish, T. N., Dircksen, H., & Gáliková, M. (2020). The influence of developmental diet on reproduction and metabolism in *Drosophila*. BMC Evolutionary Biology, 20, 1–15. 10.1186/s12862-020-01663-y

40. Kotrschal, A., Rogell, B., Bundsen, A., Svensson, B., Zajitschek, S., Brännström, I., Immler, S., Maklakov, A. A., & Kolm, N. (2013). Artificial selection on relative brain size in the guppy reveals costs and benefits of evolving a larger brain. Current Biology, 23, 168–171. 10.1016/j.cub.2012.11.058

41. Kranz, A. M., Forgan, L. G., Cole, G. L., & Endler, J. A. (2018). Light environment change induces differential expression of guppy opsins in a multi-generational evolution experiment. Evolution, 72, 1656–1676. 10.1111/evo.13519

42. Le Pape, O., & Bonhommeau, S. (2015). The food limitation hypothesis for juvenile marine fish. Fish and Fisheries, 16, 373–398. 10.1111/faf.12063

43. Lee, H. W., Ban, S., Ikada, T., & Matsuishi, T. (2003). Effect of temperature on development, growth and reproduction in the marine copepod *Pseudocalanus newmani* at satiating food condition. Journal of Plankton Research, 25, 261–271.

44. Lee, W. S., Monaghan, P., & Metcalfe, N. B. (2013). Experimental demonstration of the growth rate-lifespan trade-off. Proceedings of the Royal Society B-Biological Sciences, 280, 20122370. 10.1098/rspb.2012.2370

45. Lindholm, A. K., Head, M. L., Brooks, R. C., Rollins, L. A., Ingleby, F. C., & Zajitschek, S. R. K. (2014). Causes of male sexual trait divergence in introduced populations of guppies. Journal of Evolutionary Biology, 27, 437–448. 10.1111/jeb.12313

46. Lister, B. C., & Garcia, A. (2018). Climate-driven declines in arthropod abundance restructure a rainforest food web. Proceedings of the National Academy of Sciences of the United States of America, 115, 10397–10406. 10.1073/pnas.1722477115

47. Macartney, E. L., Crean, A. J., Nakagawa, S., & Bonduriansky, R. (2019). Effects of nutrient limitation on sperm and seminal fluid: a systematic review and meta-analysis. Biological Reviews, 94, 1722–1739. 10.1111/brv.12524

48. Marshall, D. J., Pettersen, A. K., Bode, M., & White, C. R. (2020). Developmental cost theory predicts thermal environment and vulnerability to global warming. Nature Ecology & Evolution, 4, 406–411. 10.1038/s41559-020-1114-9

49. Marshall, D. J., Pettersen, A. K., & Cameron, H. (2018). A global synthesis of offspring size variation, its eco-evolutionary causes and consequences. Functional Ecology, 32, 1436–1446. 10.1111/1365-2435.13099

50. Mattson, M. P., Allison, D. B., Fontana, L., Harvie, M., Longo, V. D., Malaisse, W. J., Mosley, M., Notterpek, L., Ravussin, E., Scheer, F. A. J. L., Seyfried, T. N., Varady, K. A., & Panda, S. (2014). Meal frequency and timing in health and disease. Proceedings of the National Academy of Sciences of the United States of America, 111, 16647–16653. 10.1073/pnas.1413965111

51. Maunder, M. N., & Piner, K. R. (2015). Contemporary fisheries stock assessment: many issues still remain. ICES Journal of Marine Science, 72, 7–18. 10.1093/icesjms/fsu015

52. Mauvais-Jarvis, F. (2024). Sex differences in energy metabolism: natural selection, mechanisms and consequences. Nature Reviews Nephrology, 20, 56–69. 10.1038/s41581-023-00781-2

53. Metcalfe, N. B., & Monaghan, P. (2001). Compensation for a bad start: grow now, pay later? Trends in Ecology & Evolution, 16, 254–260. 10.1016/S0169-5347(01)02124-3

54. Monaghan, P., & Ozanne, S. E. (2018). Somatic growth and telomere dynamics in vertebrates: relationships, mechanisms and consequences. Philosophical Transactions of the Royal Society B-Biological Sciences, 373, 20160446. 10.1098/rstb.2016.0446

55. Monteforte, S., Cattelan, S., Morosinotto, C., Pilastro, A., & Grapputo, A. (2020). Maternal predator-exposure affects offspring size at birth but not telomere length in a live-bearing fish. Ecology and Evolution, 10, 2030–2039. 10.1002/ece3.6035

56. Morbiato, E., Cattelan, S., Pilastro, A., & Grapputo, A. (2023). Sperm production is negatively associated with muscle and sperm telomere length in a species subjected to strong sperm competition. Molecular Ecology, 32, 5812–5822. 10.1111/mec.17158

57. Morrongiello, J. R., Bond, N. R., Crook, D. A., & Wong, B. B. M. (2012). Spatial variation in egg size and egg number reflects trade-offs and bet-hedging in a freshwater fish. Journal of Animal Ecology, 81, 806–817. 10.1111/j.1365-2656.2012.01961.x

58. Moura-Campos, D., Chung, M. H. J., Lawrence, E., Jennions, M. J., Head, M. L. (2024) Temperature-dependent differences in male and female life history responses to a period of food limitation during development. EcoEvoRxiv

59. Mousavi-Sabet, H., Azimi, H., Eagderi, S., Bozorgi, S., & Mahallatipour, B. (2014). Growth and morphological development of guppy *Poecilia reticulata* (Cyprinodontiformes, Poeciliidae) larvae. Poeciliid Research, 4, 24–30.

60. Munch, S. B., & Salinas, S. (2009). Latitudinal variation in lifespan within species is explained by the metabolic theory of ecology. Proceedings of the National Academy of Sciences of the United States of America, 106, 13860–13864. 10.1073/pnas.0900300106

61. Mundinger, C., Fleischer, T., Scheuerlein, A., & Kerth, G. (2022). Global warming leads to larger bats with a faster life history pace in the long-lived Bechstein’s bat (*Myotis bechsteinii*). Communications Biology, 5, 682. 10.1038/s42003-022-03611-6

62. Nakagawa, S., Kar, F., O’Dea, R. E., Pick, J. L., & Lagisz, M. (2017). Divide and conquer? Size adjustment with allometry and intermediate outcomes. BMC Biology, 15, 107. 10.1186/s12915-017-0448-5

63. Niu, J. Y., Huss, M., Vasemägi, A., & Gårdmark, A. (2023). Decades of warming alters maturation and reproductive investment in fish. Ecosphere, 14, e4381. 10.1002/ecs2.4381

64. Petitjean, Q., Jacquin, L., Riem, L., Pitout, M., Perrault, A., Cousseau, M., Laffaille, P., & Jean, S. (2021). Intraspecific variability of responses to combined metal contamination and immune challenge among wild fish populations. Environmental Pollution, 272, 16042. 10.1016/j.envpol.2020.116042

65. Ricklefs, R. E., & Cadena, C. D. (2007). Lifespan is unrelated to investment in reproduction in populations of mammals and birds in captivity. Ecology Letters, 10, 867–872. 10.1111/j.1461-0248.2007.01085.x

66. Riemer, K., Anderson-Teixeira, K. J., Smith, F. A., Harris, D. J., & Ernest, S. K. M. (2018). Body size shifts influence effects of increasing temperatures on ectotherm metabolism. Global Ecology and Biogeography, 27, 958–967. 10.1111/geb.12757

67. Roff, D. A. (1992). The Evolution of Life Histories, Theory and Analysis. Chapman and Hall.

68. Rollinson, N., & Rowe, L. (2016). The positive correlation between maternal size and offspring size: fitting pieces of a life-history puzzle. Biological Reviews, 91, 1134–1148. 10.1111/brv.12214

69. Rowiński, P. K., & Rogell, B. (2017). Environmental stress correlates with increases in both genetic and residual variances: a meta-analysis of animal studies. Evolution, 71, 1339–1351. 10.1111/evo.13201

70. Ruf, T., Fietz, J., Schlund, W., & Bieber, C. (2006). High survival in poor years: life history tactics adapted to mast seeding in the edible dormouse. Ecology, 87, 372–381. 10.1890/05-0672

71. Salinas, S., & Munch, S. B. (2012). Thermal legacies: transgenerational effects of temperature on growth in a vertebrate. Ecology Letters, 15, 159–163. 10.1111/j.1461-0248.2011.01721.x

72. Salmón, P., Nilsson, J. F., Watson, H., Bensch, S., & Isaksson, C. (2017). Selective disappearance of great tits with short telomeres in urban areas. Proceedings of the Royal Society B-Biological Sciences, 284, 20171349. 10.1098/rspb.2017.1349

73. Sandblom, E., Gräns, A., Axelsson, M., & Seth, H. (2014). Temperature acclimation rate of aerobic scope and feeding metabolism in fishes: implications in a thermally extreme future. Proceedings of the Royal Society B-Biological Sciences, 281, 20141490. 10.1098/rspb.2014.1490

74. Schulte, P. M. (2015). The effects of temperature on aerobic metabolism: towards a mechanistic understanding of the responses of ectotherms to a changing environment. Journal of Experimental Biology, 218, 1856–1866. 10.1242/jeb.118851

75. Shahjahan, R. M., Ahmed, M. J., Begum, R. A., & Rashid, M. A. (2013). Breeding biology of guppy fish, *Poecilia reticulata* (Peters, 1859) in the laboratory. Journal of the Asiatic Society of Bangladesh, 39, 259–267.

76. Sinervo, B., Méndez-de-la-Cruz, F., Miles, D. B., Heulin, B., Bastiaans, E., Cruz, M. V. S., Lara-Resendiz, R., Martínez-Méndez, N., Calderón-Espinosa, M. L., Meza-Lázaro, R. N., Gadsden, H., Avila, L. J., Morando, M., De la Riva, I. J., Sepulveda, P. V., Rocha, C. F. D., Ibargüengoytía, N., Puntriano, C. A., Massot, M., Lepetz, V., Oksanen, T. A., Chapple, D. G., Bauer, A. M., Branch, W. R., Clobert, J., & Sites, J. W. (2010). Erosion of lizard diversity by climate change and altered thermal niches. Science, 328, 894–899. 10.1126/science.1184695

77. Stearns, S. C. (1992). The Evolution of Life Histories. Oxford University Press.

78. Suikkanen, S., Pulina, S., Engström-Öst, J., Lehtiniemi, M., Lehtinen, S., & Brutemark, A. (2013). Climate change and eutrophication induced shifts in northern summer plankton communities. PLoS One, 8, e66475. 10.1371/journal.pone.0066475

79. Teder, T., & Kaasik, A. (2023). Early-life food stress hits females harder than males in insects: a meta-analysis of sex differences in environmental sensitivity. Ecology Letters, 26, 1419–1431. 10.1111/ele.14241

80. Thomas, M. K., Aranguren-Gassis, M., Kremer, C. T., Gould, M. R., Anderson, K., Klausmeier, C. A., & Litchman, E. (2017). Temperature-nutrient interactions exacerbate sensitivity to warming in phytoplankton. Global Change Biology, 23, 3269–3280. 10.1111/gcb.13641

81. Van Doorslaer, W., Stoks, R., Jeppesen, E., & De Meester, L. (2007). Adaptive microevolutionary responses to simulated global warming in *Simocephalus vetulus*: a mesocosm study. Global Change Biology, 13, 878–886. 10.1111/j.1365-2486.2007.01317.x

82. van Noordwijk, A. J., & de Jong, G. (1986). Acquisition and allocation of resources: their influence on variation in life history tactics. American Naturalist, 128, 137–142.

83. Vega-Trejo, R., Head, M. L., & Jennions, M. D. (2016). Inbreeding depression does not increase after exposure to a stressful environment: a test using compensatory growth. BMC Evolutionary Biology, 16, 68. 10.1186/s12862-016-0640-1

84. Wang, H. Y., Shen, S. F., Chen, Y. S., Kiang, Y. K., & Heino, M. (2020). Life histories determine divergent population trends for fishes under climate warming. Nature Communications, 11, 4088. 10.1038/s41467-020-17937-4

85. White, C. R., Alton, L. A., Bywater, C. L., Lombardi, E. J., & Marshall, D. J. (2022). Metabolic scaling is the product of life-history optimization. Science, 377, 834–839. 10.1126/science.abm7649

86. Wootton, H. F., Morrongiello, J. R., Schmitt, T., & Audzijonyte, A. (2022). Smaller adult fish size in warmer water is not explained by elevated metabolism. Ecology Letters, 25, 1177–1188. 10.1111/ele.13989

87. Wootton, R. J., & Smith, C. (2015). Reproductive Biology of Teleost Fishes. Wiley Blackwell.

88. Yang, J., Lu, J. H., Chen, Y., Yan, E. R., Hu, J. H., Wang, X. H., & Shen, G. C. (2020). Large underestimation of intraspecific trait variation and its improvements. Frontiers in Plant Science, 11, 53. 10.3389/fpls.2020.00053

89. Zheng, S. L., Hu, J. T., Ma, Z. J., Lindenmayer, D., & Liu, J. J. (2023). Increases in intraspecific body size variation are common among North American mammals and birds between 1880 and 2020. Nature Ecology & Evolution, 7, 347–354. 10.1038/s41559-022-01967-w

90. Zhu, Z. M., Lin, X. T., Pan, J. X., & Xu, Z. N. (2016). Effect of cyclical feeding on compensatory growth, nitrogen and phosphorus budgets in juvenile *Litopenaeus vannamei*. Aquaculture Research, 47, 283–289. 10.1111/are.12490

