## Supplementary file for "Does early-life food shortage alter the effect of elevated temperature on female life history?"

**Table S1. GLMM formulation for each measured traits. Temperature (T), early-life food availability (F), brood order (B) and adult age (A) are shown in fixed effects.**

| Trait | Error distribution | Zero-inflated | Fixed effects | Random effects |
|---|---|---|---|---|
| *Adult growth* | | | | |
| Famale size | Gaussian | No | T, F, A, $A^2$ | family ID, female ID |
| *Reproductive investment across broods* | | | | |
| Gestation time | Conway-Maxwell Poisson | No | T, F, B | family ID, female ID |
| Offspring number | Conway-Maxwell Poisson | No | T, F, B | family ID, female ID |
| Offspring size | Gaussian | No | T, F, B | family ID, female ID, brood ID |
| Variability of offspring size | Gaussian | No | T, F, B | family ID, female ID |
| *Overall fecundity* | | | | |
| Total brood number | Conway-Maxwell Poisson | Yes | T, F | family ID |
| Total offspring number | Conway-Maxwell Poisson | Yes | T, F | family ID |
| Total egg number | Gaussian | No | T, F | family ID |
| Embryo number | Conway-Maxwell Poisson | Yes | T, F | family ID |
| *Somatic investment* | | | | |
| Relative telomere length | Gaussian | No | T, F | family ID |
| Immune response | quasi-Poisson | Yes | T, F, pre-injection body tickness | family ID |
| Gut length | Gaussian | No | T, F, body size | family ID |

**Table S2. Statistical outputs of the effects of temperature, early-life food availability and age on adult growth**

**(A) Initial model including the three-way interaction among temperature, early-life food availability and age**

| Fixed effect | Estimate | *SE* | $\chi^2$ | *df* | *P* |
|---|---|---|---|---|---|
| Intercept (24°C, control) | 18.358 | 0.198 | 8625.361 | 1 | **<0.001** |
| Temperature (28°C) | 0.866 | 0.197 | 19.382 | 1 | **<0.001** |
| Food (restricted) | -0.372 | 0.194 | 3.683 | 1 | 0.055 |
| Age | 0.852 | 0.017 | 2574.955 | 1 | **<0.001** |
| $Age^2$ | -0.022 | 0.001 | 816.234 | 1 | **<0.001** |
| Temperature (28°C) * Food (restricted) | 0.605 | 0.282 | 4.590 | 1 | **0.032** |
| Temperature (28°C) * Age | -0.101 | 0.011 | 92.076 | 1 | **<0.001** |
| Food (restricted) * Age | 0.010 | 0.010 | 1.090 | 1 | 0.296 |
| Temperature (28°C) * Food (restricted) * Age | -0.034 | 0.015 | 5.249 | 1 | **0.022** |

| Random effect | Variance | *sd* | Number of groups |
|---|---|---|---|
| Family ID (intercept) | 0.452 | 0.672 | 26 |
| Female ID (intercept) | 0.635 | 0.797 | 212 |
| Residuals | 0.409 | 0.640 | |

Due to a significant three-way interaction, the interaction between temperature and early-life food availability was examined separately at each age:

**(B) Week 2 after maturation**

(i) Initial model with the interactive effect

| Fixed effect | Estimate | *SE* | $\chi^2$ | *df* | *P* |
|---|---|---|---|---|---|
| Intercept (24°C, control) | 19.773 | 0.192 | 10580.332 | 1 | **<0.001** |
| Temperature (28°C) | 0.746 | 0.205 | 13.240 | 1 | **<0.001** |
| Food (restricted) | -0.281 | 0.203 | 1.922 | 1 | 0.166 |
| Temperature (28°C) * Food (restricted) | 0.400 | 0.294 | 1.848 | 1 | 0.174 |

| Random effect | Variance | *sd* | Number of groups |
|---|---|---|---|
| Family ID (intercept) | 0.416 | 0.645 | 26 |
| Residuals | 1.059 | 1.029 | |

(ii) Final model excluding the non-significant interaction

| Fixed effect | Estimate | *SE* | $\chi^2$ | *df* | *P* |
|---|---|---|---|---|---|
| Intercept (24°C, control) | 19.684 | 0.182 | | | |
| Temperature (28°C) | 0.941 | 0.147 | 41.091 | 1 | **<0.001** |
| Food (restricted) | -0.090 | 0.146 | 0.374 | 1 | 0.541 |

| Random effect | Variance | *sd* | Number of groups |
|---|---|---|---|
| Family ID (intercept) | 0.422 | 0.650 | 26 |
| Residuals | 1.068 | 1.033 | |

(iii) Exclusion of the non-significant interaction did not significantly reduce model fit in the final model

| | *df* | AIC | BIC | Loglikelihood | Deviance | $\chi^2_1$ | *P* |
|---|---|---|---|---|---|---|---|
| Final model (ii) | 5 | 660.1 | 676.9 | -325.05 | 650.11 | | |
| Initial model (i) | 6 | 660.3 | 680.4 | -324.13 | 648.27 | 1.841 | 0.175 |

### (C) Week 6 after maturation

(i) Initial model with the interactive effect

| Fixed effect | Estimate | *SE* | $\chi^2$ | *df* | *P* |
|---|---|---|---|---|---|
| Intercept (24°C, control) | 23.066 | 0.212 | 11857.695 | 1 | **<0.001** |
| Temperature (28°C) | 0.202 | 0.207 | 0.950 | 1 | 0.330 |
| Food (restricted) | -0.424 | 0.201 | 4.472 | 1 | **0.034** |
| Temperature (28°C) * Food (restricted) | 0.528 | 0.297 | 3.167 | 1 | 0.075 |

| Random effect | Variance | *sd* | Number of groups |
|---|---|---|---|
| Family ID (intercept) | 0.624 | 0.790 | 26 |
| Residuals | 1.016 | 1.008 | |

(ii) Final model excluding the non-significant interaction

| Fixed effect | Estimate | *SE* | $\chi^2$ | *df* | *P* |
|---|---|---|---|---|---|
| Intercept (24°C, control) | 22.954 | 0.204 | | | |
| Temperature (28°C) | 0.463 | 0.147 | 9.953 | 1 | **0.002** |
| Food (restricted) | -0.180 | 0.148 | 1.486 | 1 | 0.223 |

| Random effect | Variance | *sd* | Number of groups |
|---|---|---|---|
| Family ID (intercept) | 0.634 | 0.796 | 26 |
| Residuals | 1.032 | 1.016 | |

(iii) Exclusion of the non-significant interaction did not significantly reduce model fit in the final model

| | *df* | AIC | BIC | Loglikelihood | Deviance | $\chi^2_1$ | *P* |
|---|---|---|---|---|---|---|---|
| Final model (ii) | 5 | 640.6 | 657.2 | -315.29 | 630.58 | | |
| Initial model (i) | 6 | 639.4 | 659.4 | -313.71 | 627.43 | 3.145 | 0.076 |

### (D) Week 10 after maturation

(i) Initial model with the interactive effect

| Fixed effect | Estimate | *SE* | $\chi^2$ | *df* | *P* |
|---|---|---|---|---|---|
| Intercept (24°C, control) | 24.582 | 0.189 | 16881.365 | 1 | **<0.001** |
| Temperature (28°C) | -0.013 | 0.203 | 0.004 | 1 | 0.951 |
| Food (restricted) | -0.315 | 0.194 | 2.634 | 1 | 0.105 |
| Temperature (28°C) * Food (restricted) | 0.169 | 0.291 | 0.336 | 1 | 0.562 |

| Random effect | Variance | *sd* | Number of groups |
|---|---|---|---|
| Family ID (intercept) | 0.426 | 0.653 | 26 |
| Residuals | 0.961 | 0.980 | |

(ii) Final model excluding the non-significant interaction

| Fixed effect | Estimate | *SE* | $\chi^2$ | *df* | *P* |
|---|---|---|---|---|---|
| Intercept (24°C, control) | 24.547 | 0.180 | | | |
| Temperature (28°C) | 0.071 | 0.143 | 0.246 | 1 | 0.620 |
| Food (restricted) | -0.240 | 0.144 | 2.769 | 1 | 0.096 |

| Random effect | Variance | *sd* | Number of groups |
|---|---|---|---|
| Family ID (intercept) | 0.428 | 0.654 | 26 |
| Residuals | 0.962 | 0.981 | |

(iii) Exclusion of the non-significant interaction did not significantly reduce model fit in the final model

| | *df* | AIC | BIC | Loglikelihood | Deviance | $\chi^2_1$ | *P* |
|---|---|---|---|---|---|---|---|
| Final model (ii) | 5 | 605.8 | 622.2 | -297.87 | 595.75 | | |
| Initial model (i) | 6 | 607.4 | 627.2 | -297.70 | 595.41 | 0.336 | 0.562 |

### (E) Week 14 after maturation

(i) Initial model with the interactive effect

| Fixed effect | Estimate | *SE* | $\chi^2$ | *df* | *P* |
|---|---|---|---|---|---|
| Intercept (24°C, control) | 25.834 | 0.187 | 19025.165 | 1 | **<0.001** |
| Temperature (28°C) | -0.579 | 0.208 | 7.730 | 1 | **0.005** |
| Food (restricted) | -0.181 | 0.190 | 0.900 | 1 | 0.343 |
| Temperature (28°C) * Food (restricted) | 0.049 | 0.296 | 0.027 | 1 | 0.869 |

| Random effect | Variance | *sd* | Number of groups |
|---|---|---|---|
| Family ID (intercept) | 0.427 | 0.654 | 26 |
| Residuals | 0.930 | 0.964 | |

(ii) Final model excluding the non-significant interaction

| Fixed effect | Estimate | *SE* | $\chi^2$ | *df* | *P* |
|---|---|---|---|---|---|
| Intercept (24°C, control) | 25.825 | 0.179 | | | |
| Temperature (28°C) | -0.555 | 0.146 | 14.375 | 1 | **<0.001** |
| Food (restricted) | -0.160 | 0.146 | 1.212 | 1 | 0.271 |

| Random effect | Variance | *sd* | Number of groups |
|---|---|---|---|
| Family ID (intercept) | 0.428 | 0.654 | 26 |
| Residuals | 0.930 | 0.964 | |

(iii) Exclusion of the non-significant interaction did not significantly reduce model fit in the final model

| | *df* | AIC | BIC | Loglikelihood | Deviance | $\chi^2_1$ | *P* |
|---|---|---|---|---|---|---|---|
| Final model (ii) | 5 | 571.0 | 587.2 | -280.5 | 561.01 | | |
| Initial model (i) | 6 | 573.0 | 592.5 | -280.49 | 560.98 | 0.027 | 0.869 |

### (F) Week 18 after maturation

(i) Initial model with the interactive effect

| Fixed effect | Estimate | *SE* | $\chi^2$ | *df* | *P* |
|---|---|---|---|---|---|
| Intercept (24°C, control) | 26.617 | 0.197 | 18305.938 | 1 | **<0.001** |
| Temperature (28°C) | -0.873 | 0.228 | 14.601 | 1 | **<0.001** |
| Food (restricted) | -0.199 | 0.200 | 0.989 | 1 | 0.320 |
| Temperature (28°C) * Food (restricted) | 0.046 | 0.325 | 0.020 | 1 | 0.887 |

| Random effect | Variance | *sd* | Number of groups |
|---|---|---|---|
| Family ID (intercept) | 0.463 | 0.681 | 26 |
| Residuals | 1.016 | 1.008 | |

(ii) Final model excluding the non-significant interaction

| Fixed effect | Estimate | *SE* | $\chi^2$ | *df* | *P* |
|---|---|---|---|---|---|
| Intercept (24°C, control) | 26.609 | 0.189 | | | |
| Temperature (28°C) | -0.850 | 0.162 | 27.666 | 1 | **<0.001** |
| Food (restricted) | -0.182 | 0.158 | 1.320 | 1 | 0.251 |

| Random effect | Variance | *sd* | Number of groups |
|---|---|---|---|
| Family ID (intercept) | 0.463 | 0.681 | 26 |
| Residuals | 1.016 | 1.008 | |

(iii) Exclusion of the non-significant interaction did not significantly reduce model fit in the final model

| | *df* | AIC | BIC | Loglikelihood | Deviance | $\chi^2_1$ | *P* |
|---|---|---|---|---|---|---|---|
| Final model (ii) | 5 | 546.2 | 562.1 | -268.1 | 536.2 | | |
| Initial model (i) | 6 | 548.2 | 567.2 | -268.09 | 536.18 | 0.020 | 0.887 |

**Table S3. Statistical outputs of the effects of temperature, early-life food availability, and brood order on reproductive investment across broods**

#### (A) Gestation time

(i) Initial model with the three-way interaction

| Fixed effect | Estimate | *SE* | $\chi^2$ | *df* | *P* |
|---|---|---|---|---|---|
| Intercept (24°C, control, 1st) | 3.866 | 0.044 | 7676.216 | 1 | **<0.001** |
| Temperature (28°C) | -0.323 | 0.069 | 21.784 | 1 | **<0.001** |
| Food (restricted) | -0.054 | 0.064 | 0.728 | 1 | 0.393 |
| Brood order (2nd) | -0.466 | 0.048 | 127.085 | 3 | **<0.001** |
| Brood order (3rd) | -0.457 | 0.055 | | | |
| Brood order (4th) | -0.422 | 0.083 | | | |
| Temperature (28°C) * Food (restricted) | 0.035 | 0.099 | 0.125 | 1 | 0.723 |
| Temperature (28°C) * Brood order (2nd) | 0.254 | 0.078 | 11.520 | 3 | **0.009** |
| Temperature (28°C) * Brood order (3rd) | 0.175 | 0.087 | | | |
| Temperature (28°C) * Brood order (4th) | 0.156 | 0.121 | | | |
| Food (restricted) * Brood order (2nd) | -0.007 | 0.067 | 0.866 | 3 | 0.834 |
| Food (restricted) * Brood order (3rd) | 0.058 | 0.077 | | | |
| Food (restricted) * Brood order (4th) | -0.035 | 0.127 | | | |
| Temperature (28°C) * Food (restricted) * Brood order (2nd) | -0.029 | 0.110 | 6.196 | 3 | 0.102 |
| Temperature (28°C) * Food (restricted) * Brood order (3rd) | 0.204 | 0.126 | | | |
| Temperature (28°C) * Food (restricted) * Brood order (4th) | 0.342 | 0.184 | | | |

| Random effect | Variance | *sd* | Number of groups |
|---|---|---|---|
| Family ID (intercept) | <0.001 | <0.001 | 26 |
| Female ID (intercept) | 0.063 | 0.251 | 188 |

(ii) Model with the three two-way interactions

| Fixed effect | Estimate | *SE* | $\chi^2$ | *df* | *P* |
|---|---|---|---|---|---|
| Intercept (24°C, control, 1$^{st}$) | 3.876 | 0.043 | 8100.613 | 1 | **<0.001** |
| Temperature (28°C) | -0.348 | 0.065 | 28.619 | 1 | **<0.001** |
| Food (restricted) | -0.075 | 0.061 | 1.495 | 1 | 0.221 |
| Brood order (2$^{nd}$) | -0.460 | 0.043 | 165.323 | 3 | **<0.001** |
| Brood order (3$^{rd}$) | -0.496 | 0.050 | | | |
| Brood order (4$^{th}$) | -0.494 | 0.076 | | | |
| Temperature (28°C) * Food (restricted) | 0.087 | 0.086 | 1.029 | 1 | 0.310 |
| Temperature (28°C) * Brood order (2$^{nd}$) | 0.238 | 0.056 | 30.009 | 3 | **<0.001** |
| Temperature (28°C) * Brood order (3$^{rd}$) | 0.270 | 0.064 | | | |
| Temperature (28°C) * Brood order (4$^{th}$) | 0.302 | 0.092 | | | |
| Food (restricted) * Brood order (2$^{nd}$) | -0.020 | 0.054 | 7.322 | 3 | 0.062 |
| Food (restricted) * Brood order (3$^{rd}$) | 0.135 | 0.062 | | | |
| Food (restricted) * Brood order (4$^{th}$) | 0.125 | 0.093 | | | |

| Random effect | Variance | *sd* | Number of groups |
|---|---|---|---|
| Family ID (intercept) | <0.001 | <0.001 | 26 |
| Female ID (intercept) | 0.061 | 0.248 | 188 |

(iii) Final model excluding the non-significant two-way interactions

| Fixed effect | Estimate | *SE* | $\chi^2$ | *df* | *P* |
|---|---|---|---|---|---|
| Intercept (24°C, control, 1st) | 3.846 | 0.038 | 10176.166 | 1 | **<0.001** |
| Temperature (28°C) | -0.306 | 0.050 | 37.023 | 1 | **<0.001** |
| Brood order (2nd) | -0.471 | 0.034 | 236.596 | 3 | **<0.001** |
| Brood order (3rd) | -0.432 | 0.040 | | | |
| Brood order (4th) | -0.440 | 0.065 | | | |
| Food (restricted) | -0.010 | 0.043 | 0.052 | 1 | 0.819 |
| Temperature (28°C) * Brood order (2nd) | 0.238 | 0.056 | 28.650 | 3 | **<0.001** |
| Temperature (28°C) * Brood order (3rd) | 0.263 | 0.064 | | | |
| Temperature (28°C) * Brood order (4th) | 0.298 | 0.093 | | | |

| Random effect | Variance | *sd* | Number of groups |
|---|---|---|---|
| Family ID (intercept) | <0.001 | 0.011 | 26 |
| Female ID (intercept) | 0.061 | 0.247 | 188 |

Given a significant temperature*brood order interaction, we ran pairwise comparison:

| temperature | contrast | ratio | SE | *df* | t.ratio | *P* |
|---|---|---|---|---|---|---|
| 24°C | brood 1st vs 2nd | 1.60 | 0.055 | 517 | 13.680 | **<0.001** |
| 24°C | brood 1st vs 3rd | 1.54 | 0.062 | 517 | 10.717 | **<0.001** |
| 24°C | brood 1st vs 4th | 1.55 | 0.101 | 517 | 6.758 | **<0.001** |
| 24°C | brood 2nd vs 3rd | 0.96 | 0.042 | 517 | -0.907 | 0.801 |
| 24°C | brood 2nd vs 4th | 0.97 | 0.065 | 517 | -0.457 | 0.968 |
| 24°C | brood 3rd vs 4th | 1.01 | 0.068 | 517 | 0.128 | 0.999 |
| 28°C | brood 1st vs 2nd | 1.262 | 0.057 | 517 | 5.154 | **<0.001** |
| 28°C | brood 1st vs 3rd | 1.183 | 0.061 | 517 | 3.252 | **0.007** |
| 28°C | brood 1st vs 4th | 1.153 | 0.080 | 517 | 2.069 | 0.165 |
| 28°C | brood 2nd vs 3rd | 0.938 | 0.051 | 517 | -1.189 | 0.634 |
| 28°C | brood 2nd vs 4th | 0.914 | 0.064 | 517 | -1.275 | 0.579 |
| 28°C | brood 3rd vs 4th | 0.975 | 0.070 | 517 | -0.354 | 0.985 |

| brood order | contrast | ratio | SE | *df* | t.ratio | *P* |
|---|---|---|---|---|---|---|
| 1st | 24 vs 28°C | 1.36 | 0.068 | 517 | 6.085 | **<0.001** |
| 2nd | 24 vs 28°C | 1.07 | 0.062 | 517 | 1.158 | 0.247 |
| 3rd | 24 vs 28°C | 1.04 | 0.068 | 517 | 0.646 | 0.519 |
| 4th | 24 vs 28°C | 1.01 | 0.0947 | 517 | 0.086 | 0.932 |

(iv) Model comparison

| | *df* | AIC | BIC | Loglikelihood | Deviance | $\chi^2_4$ | *P* |
|---|---|---|---|---|---|---|---|
| model (iii) | 12 | 3970.4 | 4021.6 | -1973.2 | 3946.4 | | |
| model (ii) | 16 | 3970.1 | 4038.4 | -1969 | 3938.1 | 8.283 | 0.082 |

(v) An additional model testing the effect of absolute body length

| Fixed effect | Estimate | *SE* | $\chi^2$ | *df* | *P* |
|---|---|---|---|---|---|
| Intercept (24°C, control, 1st) | 3.959 | 0.039 | 10478.070 | 1 | **<0.001** |
| Temperature (28°C) | -0.241 | 0.039 | 37.753 | 1 | **<0.001** |
| Brood order (2nd) | -0.713 | 0.034 | 556.260 | 3 | **<0.001** |
| Brood order (3rd) | -0.845 | 0.043 | | | |
| Brood order (4th) | -0.957 | 0.066 | | | |
| Food (restricted) | 0.035 | 0.033 | 1.116 | 1 | 0.291 |
| Female size (standardized) | 0.279 | 0.019 | 208.679 | 1 | **<0.001** |
| Temperature (28°C) * Brood order (2nd) | 0.303 | 0.048 | 69.106 | 3 | **<0.001** |
| Temperature (28°C) * Brood order (3rd) | 0.371 | 0.056 | | | |
| Temperature (28°C) * Brood order (4th) | 0.419 | 0.081 | | | |

| Random effect | Variance | *sd* | Number of groups |
|---|---|---|---|
| Family ID (intercept) | 0.013 | 0.115 | 26 |
| Female ID (intercept) | 0.029 | 0.170 | 188 |

Given a significant temperature*brood order interaction, we ran pairwise comparison:

| temperature | contrast | ratio | SE | *df* | t.ratio | *P* |
|---|---|---|---|---|---|---|
| 24°C | brood 1st vs 2nd | 2.040 | 0.069 | 513 | 21.208 | **<0.001** |
| 24°C | brood 1st vs 3rd | 2.330 | 0.101 | 513 | 19.579 | **<0.001** |
| 24°C | brood 1st vs 4th | 2.600 | 0.173 | 513 | 14.438 | **<0.001** |
| 24°C | brood 2nd vs 3rd | 1.140 | 0.043 | 513 | 3.464 | **0.003** |
| 24°C | brood 2nd vs 4th | 1.280 | 0.078 | 513 | 4.003 | **<0.001** |
| 24°C | brood 3rd vs 4th | 1.120 | 0.067 | 513 | 1.876 | 0.240 |
| 28°C | brood 1st vs 2nd | 1.510 | 0.060 | 513 | 10.370 | **<0.001** |
| 28°C | brood 1st vs 3rd | 1.610 | 0.076 | 513 | 10.040 | **<0.001** |
| 28°C | brood 1st vs 4th | 1.710 | 0.107 | 513 | 8.600 | **<0.001** |
| 28°C | brood 2nd vs 3rd | 1.070 | 0.050 | 513 | 1.374 | 0.516 |
| 28°C | brood 2nd vs 4th | 1.140 | 0.069 | 513 | 2.096 | 0.156 |
| 28°C | brood 3rd vs 4th | 1.070 | 0.065 | 513 | 1.041 | 0.726 |

| brood order | contrast | ratio | SE | *df* | t.ratio | *P* |
|---|---|---|---|---|---|---|
| 1st | 24 vs 28°C | 1.272 | 0.050 | 513 | 6.144 | **<0.001** |
| 2nd | 24 vs 28°C | 0.939 | 0.044 | 513 | -1.332 | 0.184 |
| 3rd | 24 vs 28°C | 0.878 | 0.047 | 513 | -2.426 | **0.016** |
| 4th | 24 vs 28°C | 0.837 | 0.067 | 513 | -2.233 | **0.026** |

### (B) Offspring number (per brood)

#### (i) Initial model with the three-way interaction

| Fixed effect | Estimate | *SE* | $\chi^2$ | *df* | *P* |
|---|---|---|---|---|---|
| Intercept (24°C, control, 1st) | 2.301 | 0.067 | 1189.456 | 1 | **<0.001** |
| Temperature (28°C) | -0.353 | 0.105 | 11.323 | 1 | **0.001** |
| Food (restricted) | -0.024 | 0.092 | 0.069 | 1 | 0.792 |
| Brood order (2nd) | 0.258 | 0.080 | 14.931 | 3 | **0.002** |
| Brood order (3rd) | 0.294 | 0.085 | | | |
| Brood order (4th) | 0.205 | 0.121 | | | |
| Temperature (28°C) * Food (restricted) | -0.059 | 0.152 | 0.150 | 1 | 0.698 |
| Temperature (28°C) * Brood order (2nd) | -0.125 | 0.139 | 2.144 | 3 | 0.543 |
| Temperature (28°C) * Brood order (3rd) | -0.089 | 0.145 | | | |
| Temperature (28°C) * Brood order (4th) | -0.277 | 0.199 | | | |
| Food (restricted) * Brood order (2nd) | 0.012 | 0.114 | 2.768 | 3 | 0.429 |
| Food (restricted) * Brood order (3rd) | 0.091 | 0.121 | | | |
| Food (restricted) * Brood order (4th) | 0.264 | 0.174 | | | |
| Temperature (28°C) * Food (restricted) * Brood order (2nd) | 0.027 | 0.198 | 5.067 | 3 | 0.167 |
| Temperature (28°C) * Food (restricted) * Brood order (3rd) | -0.328 | 0.221 | | | |
| Temperature (28°C) * Food (restricted) * Brood order (4th) | -0.502 | 0.314 | | | |

| Random effect | Variance | *sd* | Number of groups |
|---|---|---|---|
| Family ID (intercept) | 0.008 | 0.087 | 26 |
| Female ID (intercept) | 0.033 | 0.181 | 188 |

(ii) Model with the three two-way interactions

| Fixed effect | Estimate | *SE* | $\chi^2$ | *df* | *P* |
|---|---|---|---|---|---|
| Intercept (24°C, control, 1st) | 2.283 | 0.064 | 1263.539 | 1 | **<0.001** |
| Temperature (28°C) | -0.302 | 0.090 | 11.210 | 1 | **0.001** |
| Food (restricted) | 0.014 | 0.083 | 0.030 | 1 | 0.863 |
| Brood order (2nd) | 0.256 | 0.074 | 21.766 | 3 | **<0.001** |
| Brood order (3rd) | 0.342 | 0.078 | | | |
| Brood order (4th) | 0.281 | 0.111 | | | |
| Temperature (28°C) * Food (restricted) | -0.166 | 0.103 | 2.601 | 1 | 0.107 |
| Temperature (28°C) * Brood order (2nd) | -0.110 | 0.099 | 11.398 | 3 | **0.010** |
| Temperature (28°C) * Brood order (3rd) | -0.228 | 0.109 | | | |
| Temperature (28°C) * Brood order (4th) | -0.482 | 0.153 | | | |
| Food (restricted) * Brood order (2nd) | 0.014 | 0.093 | 0.655 | 3 | 0.884 |
| Food (restricted) * Brood order (3rd) | -0.006 | 0.101 | | | |
| Food (restricted) * Brood order (4th) | 0.107 | 0.144 | | | |

| Random effect | Variance | *sd* | Number of groups |
|---|---|---|---|
| Family ID (intercept) | 0.007 | 0.083 | 26 |
| Female ID (intercept) | 0.034 | 0.185 | 188 |

(iii) Final model excluding the non-significant two-way interactions

| Fixed effect | Estimate | *SE* | $\chi^2$ | *df* | *P* |
|---|---|---|---|---|---|
| Intercept (24°C, control, 1st) | 2.303 | 0.056 | 1699.880 | 1 | **<0.001** |
| Temperature (28°C) | -0.382 | 0.076 | 25.364 | 1 | **<0.001** |
| Brood order (2nd) | 0.264 | 0.057 | 37.623 | 3 | **<0.001** |
| Brood order (3rd) | 0.339 | 0.061 | | | |
| Brood order (4th) | 0.330 | 0.087 | | | |
| Food (restricted) | -0.032 | 0.049 | 0.440 | 1 | 0.507 |
| Temperature (28°C) * Brood order (2nd) | -0.113 | 0.099 | 11.318 | 3 | **0.010** |
| Temperature (28°C) * Brood order (3rd) | -0.222 | 0.109 | | | |
| Temperature (28°C) * Brood order (4th) | -0.483 | 0.153 | | | |

| Random effect | Variance | *sd* | Number of groups |
|---|---|---|---|
| Family ID (intercept) | 0.008 | 0.090 | 26 |
| Female ID (intercept) | 0.036 | 0.190 | 188 |

Given a significant temperature*brood order interaction, we ran pairwise comparison:

| temperature | contrast | ratio | SE | *df* | t.ratio | *P* |
|---|---|---|---|---|---|---|
| 24°C | brood 1st vs 2nd | 0.768 | 0.044 | 517 | -4.636 | **<0.001** |
| 24°C | brood 1st vs 3rd | 0.712 | 0.043 | 517 | -5.585 | **<0.001** |
| 24°C | brood 1st vs 4th | 0.719 | 0.063 | 517 | -3.785 | **0.001** |
| 24°C | brood 2nd vs 3rd | 0.928 | 0.054 | 517 | -1.297 | 0.565 |
| 24°C | brood 2nd vs 4th | 0.937 | 0.080 | 517 | -0.767 | 0.869 |
| 24°C | brood 3rd vs 4th | 1.010 | 0.088 | 517 | 0.112 | 0.999 |
| 28°C | brood 1st vs 2nd | 0.859 | 0.070 | 517 | -1.863 | 0.245 |
| 28°C | brood 1st vs 3rd | 0.889 | 0.081 | 517 | -1.298 | 0.565 |
| 28°C | brood 1st vs 4th | 1.166 | 0.146 | 517 | 1.224 | 0.612 |
| 28°C | brood 2nd vs 3rd | 1.035 | 0.0943 | 517 | 0.372 | 0.982 |
| 28°C | brood 2nd vs 4th | 1.357 | 0.1708 | 517 | 2.422 | 0.074 |
| 28°C | brood 3rd vs 4th | 1.311 | 0.171 | 517 | 2.078 | 0.162 |

| brood order | contrast | ratio | SE | *df* | t.ratio | *P* |
|---|---|---|---|---|---|---|
| 1st | 24 vs 28°C | 1.470 | 0.111 | 517 | 5.036 | **<0.001** |
| 2nd | 24 vs 28°C | 1.640 | 0.125 | 517 | 6.500 | **<0.001** |
| 3rd | 24 vs 28°C | 1.830 | 0.162 | 517 | 6.830 | **<0.001** |
| 4th | 24 vs 28°C | 2.380 | 0.330 | 517 | 6.231 | **<0.001** |

### (iv) Model comparison

| | *df* | AIC | BIC | Loglikelihood | Deviance | $\chi^2_4$ | *P* |
|---|---|---|---|---|---|---|---|
| model (iii) | 12 | 3114.1 | 3165.4 | -1545.1 | 3090.1 | | |
| model (ii) | 16 | 3118.9 | 3187.2 | -1543.5 | 3086.9 | 3.237 | 0.519 |

(v) An additional model testing the effect of absolute body length

| Fixed effect | Estimate | *SE* | $\chi^2$ | *df* | *P* |
|---|---|---|---|---|---|
| Intercept (24°C, control, 1st) | 2.369 | 0.057 | 1750.826 | 1 | **<0.001** |
| Temperature (28°C) | -0.351 | 0.074 | 22.547 | 1 | **<0.001** |
| Brood order (2nd) | 0.143 | 0.062 | 7.836 | 3 | **0.0495** |
| Brood order (3rd) | 0.157 | 0.071 | | | |
| Brood order (4th) | 0.036 | 0.103 | | | |
| Food (restricted) | -0.022 | 0.046 | 0.228 | 1 | 0.633 |
| Female size (standardized) | 0.136 | 0.028 | 23.162 | 1 | **<0.001** |
| Temperature (28°C) * Brood order (2nd) | -0.074 | 0.099 | 6.788 | 3 | 0.079 |
| Temperature (28°C) * Brood order (3rd) | -0.182 | 0.108 | | | |
| Temperature (28°C) * Brood order (4th) | -0.365 | 0.154 | | | |

| Random effect | Variance | *sd* | Number of groups |
|---|---|---|---|
| Family ID (intercept) | 0.013 | 0.114 | 26 |
| Female ID (intercept) | 0.022 | 0.149 | 188 |

(vi) Final model for the effect of absolut body length (i.e., removing the non-significant interaction)

| Fixed effect | Estimate | *SE* | $\chi^2$ | *df* | *P* |
|---|---|---|---|---|---|
| Intercept (24°C, control, 1st) | 2.406 | 0.053 | | | |
| Temperature (28°C) | -0.448 | 0.049 | 83.762 | 1 | **<0.001** |
| Brood order (2nd) | 0.111 | 0.051 | 12.617 | 3 | **0.006** |
| Brood order (3rd) | 0.091 | 0.061 | | | |
| Brood order (4th) | -0.107 | 0.086 | | | |
| Food (restricted) | -0.019 | 0.046 | 0.179 | 1 | 0.672 |
| Female size (standardized) | 0.143 | 0.028 | 25.578 | 1 | **<0.001** |

| Random effect | Variance | *sd* | Number of groups |
|---|---|---|---|
| Family ID (intercept) | 0.013 | 0.114 | 26 |
| Female ID (intercept) | 0.022 | 0.148 | 188 |

Given a significant effect of brood order, we ran pairwise comparsion:

| contrast | ratio | SE | *df* | t.ratio | *P* |
|---|---|---|---|---|---|
| brood 1st vs 2nd | 0.895 | 0.046 | 516 | -2.161 | 0.136 |
| brood 1st vs 3rd | 0.913 | 0.056 | 516 | -1.484 | 0.448 |
| brood 1st vs 4th | 1.113 | 0.096 | 516 | 1.241 | 0.601 |
| brood 2nd vs 3rd | 1.021 | 0.052 | 516 | 0.407 | 0.977 |
| brood 2nd vs 4th | 1.244 | 0.095 | 516 | 2.847 | **0.024** |
| brood 3rd vs 4th | 1.218 | 0.092 | 516 | 2.627 | **0.044** |

### (C) Offspring size

(i) Initial model with the three-way interaction

| **Fixed effect** | Estimate | *SE* | $\chi^2$ | *df* | *P* |
|---|---|---|---|---|---|
| Intercept (24°C, control, 1st) | 6.468 | 0.042 | 23784.964 | 1 | **<0.001** |
| Temperature (28°C) | -0.234 | 0.055 | 18.035 | 1 | **<0.001** |
| Food (restricted) | -0.042 | 0.051 | 0.670 | 1 | 0.413 |
| Brood order (2nd) | 0.189 | 0.040 | 47.892 | 3 | **<0.001** |
| Brood order (3rd) | 0.261 | 0.044 | | | |
| Brood order (4th) | 0.317 | 0.063 | | | |
| Temperature (28°C) * Food (restricted) | 0.073 | 0.079 | 0.853 | 1 | 0.356 |
| Temperature (28°C) * Brood order (2nd) | -0.040 | 0.066 | 2.342 | 3 | 0.505 |
| Temperature (28°C) * Brood order (3rd) | -0.051 | 0.070 | | | |
| Temperature (28°C) * Brood order (4th) | -0.143 | 0.095 | | | |
| Food (restricted) * Brood order (2nd) | 0.011 | 0.058 | 1.967 | 3 | 0.579 |
| Food (restricted) * Brood order (3rd) | 0.028 | 0.063 | | | |
| Food (restricted) * Brood order (4th) | 0.137 | 0.100 | | | |
| Temperature (28°C) * Food (restricted) * Brood order (2nd) | -0.058 | 0.093 | 2.407 | 3 | 0.492 |
| Temperature (28°C) * Food (restricted) * Brood order (3rd) | -0.119 | 0.104 | | | |
| Temperature (28°C) * Food (restricted) * Brood order (4th) | 0.089 | 0.150 | | | |

| Random effect | Variance | *sd* | Number of groups |
|---|---|---|---|
| Family ID (intercept) | 0.012 | 0.110 | 26 |
| Female ID (intercept) | 0.022 | 0.149 | 188 |
| Brood ID (intercept) | 0.031 | 0.177 | 524 |
| Residual | 0.091 | 0.302 | |

(ii) Model with the three two-way interactions

| **Fixed effect** | Estimate | *SE* | $\chi^2$ | *df* | *P* |
|---|---|---|---|---|---|
| Intercept (24°C, control, 1st) | 6.461 | 0.041 | 25008.752 | 1 | **<0.001** |
| Temperature (28°C) | -0.216 | 0.050 | 18.733 | 1 | **<0.001** |
| Food (restricted) | -0.026 | 0.047 | 0.311 | 1 | 0.577 |
| Brood order (2nd) | 0.200 | 0.036 | 63.130 | 3 | **<0.001** |
| Brood order (3rd) | 0.282 | 0.040 | | | |
| Brood order (4th) | 0.302 | 0.058 | | | |
| Temperature (28°C) * Food (restricted) | 0.036 | 0.062 | 0.332 | 1 | 0.564 |
| Temperature (28°C) * Brood order (2nd) | -0.069 | 0.047 | 5.200 | 3 | 0.158 |
| Temperature (28°C) * Brood order (3rd) | -0.104 | 0.052 | | | |
| Temperature (28°C) * Brood order (4th) | -0.114 | 0.074 | | | |
| Food (restricted) * Brood order (2nd) | -0.012 | 0.045 | 7.531 | 3 | 0.057 |
| Food (restricted) * Brood order (3rd) | -0.016 | 0.050 | | | |
| Food (restricted) * Brood order (4th) | 0.180 | 0.075 | | | |

| Random effect | Variance | *sd* | Number of groups |
|---|---|---|---|
| Family ID (intercept) | 0.012 | 0.109 | 26 |
| Female ID (intercept) | 0.022 | 0.149 | 188 |
| Brood ID (intercept) | 0.031 | 0.177 | 524 |
| Residual | 0.091 | 0.302 | |

(iii) Final model excluding the non-significant two-way interactions

| **Fixed effect** | Estimate | *SE* | $\chi^2$ | *df* | *P* |
|---|---|---|---|---|---|
| Intercept (24°C, control, 1$^{st}$) | 6.474 | 0.036 | | | |
| Temperature (28°C) | -0.250 | 0.030 | 70.359 | 1 | **<0.001** |
| Food (restricted) | -0.006 | 0.030 | 0.039 | 1 | 0.844 |
| Brood order (2$^{nd}$) | 0.166 | 0.023 | 119.051 | 3 | **<0.001** |
| Brood order (3$^{rd}$) | 0.232 | 0.026 | | | |
| Brood order (4$^{th}$) | 0.317 | 0.038 | | | |

| Random effect | Variance | *sd* | Number of groups |
|---|---|---|---|
| Family ID (intercept) | 0.012 | 0.111 | 26 |
| Female ID (intercept) | 0.020 | 0.141 | 188 |
| Brood ID (intercept) | 0.034 | 0.184 | 524 |
| Residual | 0.091 | 0.302 | |

Given a significant effect of brood order, we ran pairwise comparsion:

| contrast | estimate | SE | *df* | t.ratio | *P* |
|---|---|---|---|---|---|
| brood 1$^{st}$ vs 2$^{nd}$ | -0.166 | 0.023 | 5348 | -7.118 | **<0.001** |
| brood 1$^{st}$ vs 3$^{rd}$ | -0.232 | 0.026 | 5348 | -9.019 | **<0.001** |
| brood 1$^{st}$ vs 4$^{th}$ | -0.318 | 0.038 | 5348 | -8.321 | **<0.001** |
| brood 2$^{nd}$ vs 3$^{rd}$ | -0.066 | 0.026 | 5348 | -2.581 | **0.049** |
| brood 2$^{nd}$ vs 4$^{th}$ | -0.151 | 0.038 | 5348 | -3.979 | **<0.001** |
| brood 3$^{rd}$ vs 4$^{th}$ | -0.085 | 0.039 | 5348 | -2.209 | 0.121 |

(iv) Model comparison

| | *df* | AIC | BIC | Loglikelihood | Deviance | $\chi^2_7$ | *P* |
|---|---|---|---|---|---|---|---|
| model (iii) | 10 | 3332.9 | 3398.8 | -1656.5 | 3312.9 | | |
| model (ii) | 17 | 3334.3 | 3446.2 | -1650.1 | 3300.3 | 12.661 | 0.081 |

(v) An additional model testing the effect of absolute body length

| **Fixed effect** | Estimate | *SE* | $\chi^2$ | *df* | *P* |
|---|---|---|---|---|---|
| Intercept (24°C, control, $1^{st}$) | 6.517 | 0.035 | | | |
| Temperature (28°C) | -0.219 | 0.030 | 53.949 | 1 | **<0.001** |
| Food (restricted) | 0.007 | 0.029 | 0.058 | 1 | 0.809 |
| Brood order ($2^{nd}$) | 0.103 | 0.026 | 20.988 | 3 | **<0.001** |
| Brood order ($3^{rd}$) | 0.127 | 0.033 | | | |
| Brood order ($4^{th}$) | 0.182 | 0.046 | | | |
| Female size (standardized) | 0.084 | 0.018 | 22.525 | 1 | **<0.001** |

| Random effect | Variance | *sd* | Number of groups |
|---|---|---|---|
| Family ID (intercept) | 0.009 | 0.097 | 26 |
| Female ID (intercept) | 0.019 | 0.138 | 188 |
| Brood ID (intercept) | 0.032 | 0.179 | 524 |
| Residual | 0.091 | 0.302 | |

Given a significant effect of brood order, we ran pairwise comparsion:

| contrast | estimate | SE | *df* | t.ratio | *P* |
|---|---|---|---|---|---|
| brood $1^{st}$ vs $2^{nd}$ | -0.103 | 0.026 | 5347 | -3.899 | **0.001** |
| brood $1^{st}$ vs $3^{rd}$ | -0.127 | 0.033 | 5347 | -3.797 | **0.001** |
| brood $1^{st}$ vs $4^{th}$ | -0.182 | 0.046 | 5347 | -3.919 | **0.001** |
| brood $2^{nd}$ vs $3^{rd}$ | -0.024 | 0.027 | 5347 | -0.906 | 0.802 |
| brood $2^{nd}$ vs $4^{th}$ | -0.079 | 0.040 | 5347 | -1.979 | 0.196 |
| brood $3^{rd}$ vs $4^{th}$ | -0.055 | 0.038 | 5347 | -1.436 | 0.477 |

### (D) Within-brood variation in offspring size

#### (i) Initial model with the three-way interaction

| Fixed effect | Estimate | *SE* | $\chi^2$ | *df* | *P* |
|---|---|---|---|---|---|
| Intercept (24°C, control, 1st) | 0.023 | 0.001 | 247.829 | 1 | **<0.001** |
| Temperature (28°C) | 0.006 | 0.002 | 7.964 | 1 | **0.005** |
| Food (restricted) | 0.001 | 0.002 | 0.130 | 1 | 0.719 |
| Brood order (2nd) | -0.001 | 0.002 | 1.696 | 3 | 0.638 |
| Brood order (3rd) | -0.001 | 0.002 | | | |
| Brood order (4th) | -0.004 | 0.003 | | | |
| Temperature (28°C) * Food (restricted) | -0.005 | 0.003 | 2.825 | 1 | 0.093 |
| Temperature (28°C) * Brood order (2nd) | -0.003 | 0.003 | 7.522 | 3 | 0.057 |
| Temperature (28°C) * Brood order (3rd) | -0.009 | 0.003 | | | |
| Temperature (28°C) * Brood order (4th) | -0.006 | 0.004 | | | |
| Food (restricted) * Brood order (2nd) | 0.000 | 0.003 | 0.303 | 3 | 0.959 |
| Food (restricted) * Brood order (3rd) | 0.002 | 0.003 | | | |
| Food (restricted) * Brood order (4th) | 0.000 | 0.005 | | | |
| Temperature (28°C) * Food (restricted) * Brood order (2nd) | 0.005 | 0.005 | 1.393 | 3 | 0.707 |
| Temperature (28°C) * Food (restricted) * Brood order (3rd) | 0.005 | 0.005 | | | |
| Temperature (28°C) * Food (restricted) * Brood order (4th) | 0.001 | 0.007 | | | |

| Random effect | Variance | *sd* | Number of groups |
|---|---|---|---|
| Family ID (intercept) | <0.001 | 0.002 | 26 |
| Female ID (intercept) | <0.001 | 0.003 | 181 |
| Residual | <0.001 | 0.010 | |

(ii) Model with the three two-way interactions

| Fixed effect | Estimate | *SE* | $\chi^2$ | *df* | *P* |
|---|---|---|---|---|---|
| Intercept (24°C, control, 1st) | 0.024 | 0.001 | 288.898 | 1 | **<0.001** |
| Temperature (28°C) | 0.005 | 0.002 | 7.007 | 1 | **0.008** |
| Food (restricted) | 0.000 | 0.002 | 0.041 | 1 | 0.839 |
| Brood order (2nd) | -0.002 | 0.002 | 2.980 | 3 | 0.395 |
| Brood order (3rd) | -0.002 | 0.002 | | | |
| Brood order (4th) | -0.004 | 0.003 | | | |
| Temperature (28°C) * Food (restricted) | -0.003 | 0.002 | 1.762 | 1 | 0.184 |
| Temperature (28°C) * Brood order (2nd) | -0.001 | 0.002 | 8.861 | 3 | **0.031** |
| Temperature (28°C) * Brood order (3rd) | -0.007 | 0.002 | | | |
| Temperature (28°C) * Brood order (4th) | -0.005 | 0.003 | | | |
| Food (restricted) * Brood order (2nd) | 0.002 | 0.002 | 2.219 | 3 | 0.528 |
| Food (restricted) * Brood order (3rd) | 0.003 | 0.002 | | | |
| Food (restricted) * Brood order (4th) | 0.000 | 0.004 | | | |

| Random effect | Variance | *sd* | Number of groups |
|---|---|---|---|
| Family ID (intercept) | <0.001 | 0.002 | 26 |
| Female ID (intercept) | <0.001 | 0.003 | 181 |
| Residual | <0.001 | 0.010 | |

(iii) Final model excluding the non-significant two-way interactions

| Fixed effect | Estimate | *SE* | $\chi^2$ | *df* | *P* |
|---|---|---|---|---|---|
| Intercept (24°C, control, 1st) | 0.024 | 0.001 | 376.675 | 1 | **<0.001** |
| Temperature (28°C) | 0.004 | 0.002 | 5.114 | 1 | **0.024** |
| Brood order (2nd) | -0.001 | 0.001 | 3.208 | 3 | 0.361 |
| Brood order (3rd) | 0.000 | 0.002 | | | |
| Brood order (4th) | -0.004 | 0.002 | | | |
| Food (restricted) | 0.000 | 0.001 | 0.001 | 1 | 0.975 |
| Temperature (28°C) * Brood order (2nd) | -0.001 | 0.002 | 8.895 | 3 | **0.031** |
| Temperature (28°C) * Brood order (3rd) | -0.007 | 0.002 | | | |
| Temperature (28°C) * Brood order (4th) | -0.004 | 0.003 | | | |

| Random effect | Variance | *sd* | Number of groups |
|---|---|---|---|
| Family ID (intercept) | <0.001 | 0.002 | 26 |
| Female ID (intercept) | <0.001 | 0.003 | 181 |
| Residual | <0.001 | 0.010 | |

Given a significant temperature*brood order interaction, we ran pairwise comparison:

| temperature | contrast | estimate | SE | *df* | t.ratio | *P* |
|---|---|---|---|---|---|---|
| 24°C | brood 1$^{st}$ vs 2$^{nd}$ | 0.001 | 0.001 | 445 | 0.861 | 0.825 |
| 24°C | brood 1$^{st}$ vs 3$^{rd}$ | 0.000 | 0.002 | 445 | 0.210 | 0.997 |
| 24°C | brood 1$^{st}$ vs 4$^{th}$ | 0.004 | 0.002 | 445 | 1.684 | 0.333 |
| 24°C | brood 2$^{nd}$ vs 3$^{rd}$ | -0.001 | 0.002 | 445 | -0.592 | 0.935 |
| 24°C | brood 2$^{nd}$ vs 4$^{th}$ | 0.003 | 0.002 | 445 | 1.159 | 0.653 |
| 24°C | brood 3$^{rd}$ vs 4$^{th}$ | 0.004 | 0.002 | 445 | 1.507 | 0.434 |
| 28°C | brood 1$^{st}$ vs 2$^{nd}$ | 0.002 | 0.002 | 445 | 1.346 | 0.534 |
| 28°C | brood 1$^{st}$ vs 3$^{rd}$ | 0.007 | 0.002 | 445 | 3.752 | **0.001** |
| 28°C | brood 1$^{st}$ vs 4$^{th}$ | 0.008 | 0.003 | 445 | 3.163 | **0.009** |
| 28°C | brood 2$^{nd}$ vs 3$^{rd}$ | 0.005 | 0.002 | 445 | 2.397 | 0.079 |
| 28°C | brood 2$^{nd}$ vs 4$^{th}$ | 0.006 | 0.003 | 445 | 2.191 | 0.127 |
| 28°C | brood 3$^{rd}$ vs 4$^{th}$ | 0.001 | 0.003 | 445 | 0.386 | 0.980 |

| brood order | contrast | estimate | SE | *df* | t.ratio | *P* |
|---|---|---|---|---|---|---|
| 1$^{st}$ | 24 vs 28°C | -0.004 | 0.002 | 445 | -2.261 | **0.024** |
| 2$^{nd}$ | 24 vs 28°C | -0.002 | 0.002 | 445 | -1.416 | 0.157 |
| 3$^{rd}$ | 24 vs 28°C | 0.003 | 0.002 | 445 | 1.679 | 0.094 |
| 4$^{th}$ | 24 vs 28°C | 0.001 | 0.003 | 445 | 0.253 | 0.800 |

(iv) Model comparison

| | *df* | AIC | BIC | Loglikelihood | Deviance | $\chi^2_4$ | *P* |
|---|---|---|---|---|---|---|---|
| model (iii) | 12 | -2889.0 | -2839.5 | 1456.5 | -2913.0 | | |
| model (ii) | 16 | -2885.2 | -2819.2 | 1458.6 | -2917.2 | 4.215 | 0.378 |

(v) An additional model testing the effect of absolute body length

| Fixed effect | Estimate | *SE* | $\chi^2$ | *df* | *P* |
|---|---|---|---|---|---|
| Intercept (24°C, control, 1st) | 0.023 | 0.001 | 352.420 | 1 | **<0.001** |
| Temperature (28°C) | 0.004 | 0.002 | 4.984 | 1 | **0.026** |
| Brood order (2nd) | -0.001 | 0.002 | 2.501 | 3 | 0.475 |
| Brood order (3rd) | 0.000 | 0.002 | | | |
| Brood order (4th) | -0.003 | 0.003 | | | |
| Food (restricted) | 0.000 | 0.001 | 0.000 | 1 | 0.997 |
| Female size (standardized) | 0.000 | 0.001 | 0.180 | 1 | 0.672 |
| Temperature (28°C) * Brood order (2nd) | -0.001 | 0.002 | 9.054 | 3 | **0.029** |
| Temperature (28°C) * Brood order (3rd) | -0.007 | 0.002 | | | |
| Temperature (28°C) * Brood order (4th) | -0.005 | 0.003 | | | |

| Random effect | Variance | *sd* | Number of groups |
|---|---|---|---|
| Family ID (intercept) | <0.001 | 0.002 | 26 |
| Female ID (intercept) | <0.001 | 0.003 | 181 |
| Residual | <0.001 | 0.010 | |

Given a significant temperature*brood order interaction, we ran pairwise comparison:

| temperature | contrast | estimate | SE | *df* | t.ratio | *P* |
|---|---|---|---|---|---|---|
| 24°C | brood 1st vs 2nd | 0.001 | 0.002 | 444 | 0.65 | 0.916 |
| 24°C | brood 1st vs 3rd | 0.000 | 0.002 | 444 | -0.024 | 1.000 |
| 24°C | brood 1st vs 4th | 0.003 | 0.003 | 444 | 1.342 | 0.537 |
| 24°C | brood 2nd vs 3rd | -0.001 | 0.002 | 444 | -0.666 | 0.910 |
| 24°C | brood 2nd vs 4th | 0.002 | 0.002 | 444 | 1.032 | 0.731 |
| 24°C | brood 3rd vs 4th | 0.003 | 0.002 | 444 | 1.461 | 0.462 |
| 28°C | brood 1st vs 2nd | 0.002 | 0.002 | 444 | 1.247 | 0.597 |
| 28°C | brood 1st vs 3rd | 0.007 | 0.002 | 444 | 3.45 | **0.003** |
| 28°C | brood 1st vs 4th | 0.008 | 0.003 | 444 | 2.911 | **0.020** |
| 28°C | brood 2nd vs 3rd | 0.005 | 0.002 | 444 | 2.317 | 0.096 |
| 28°C | brood 2nd vs 4th | 0.006 | 0.003 | 444 | 2.092 | 0.157 |
| 28°C | brood 3rd vs 4th | 0.001 | 0.003 | 444 | 0.359 | 0.984 |

| brood order | contrast | estimate | SE | *df* | t.ratio | *P* |
|---|---|---|---|---|---|---|
| 1st | 24 vs 28°C | -0.004 | 0.002 | 444 | -2.232 | **0.026** |
| 2nd | 24 vs 28°C | -0.002 | 0.002 | 444 | -1.319 | 0.188 |
| 3rd | 24 vs 28°C | 0.003 | 0.002 | 444 | 1.726 | 0.085 |
| 4th | 24 vs 28°C | 0.001 | 0.003 | 444 | 0.304 | 0.761 |

**Table S4. Effects of temperature and early-life food availability on overall fecundity**

**(A) Total brood number**

(i) Initial model with the interactive effect

Zero-inflation part

| Fixed effect | Estimate | *SE* | $\chi^2$ | *df* | *P* |
|---|---|---|---|---|---|
| Intercept (24°C, control) | -2.992 | 0.642 | 21.686 | 1 | **<0.001** |
| Temperature (28°C) | 1.767 | 0.701 | 6.350 | 1 | **0.012** |
| Food (restricted) | -0.355 | 0.981 | 0.131 | 1 | 0.717 |
| Temperature (28°C) * Food (restricted) | 0.043 | 1.096 | 0.002 | 1 | 0.969 |

| Random effect | Variance | *sd* | Number of groups |
|---|---|---|---|
| Family ID (intercept) | 0.121 | 0.348 | 26 |

Conditional part

| Fixed effect | Estimate | *SE* | $\chi^2$ | *df* | *P* |
|---|---|---|---|---|---|
| Intercept (24°C, control) | 1.059 | 0.049 | 460.720 | 1 | **<0.001** |
| Temperature (28°C) | -0.003 | 0.075 | 0.002 | 1 | 0.969 |
| Food (restricted) | 0.018 | 0.071 | 0.062 | 1 | 0.803 |
| Temperature (28°C) * Food (restricted) | -0.115 | 0.108 | 1.132 | 1 | 0.287 |

| Random effect | Variance | *sd* | Number of groups |
|---|---|---|---|
| Family ID (intercept) | <0.001 | <0.001 | 26 |

(ii) Final model excluding the non-significant interaction

Zero-inflation part

| Fixed effect | Estimate | *SE* | $\chi^2$ | *df* | *P* |
|---|---|---|---|---|---|
| Intercept (24°C, control) | -3.010 | 0.538 | | | |
| Temperature (28°C) | 1.787 | 0.540 | 10.961 | 1 | **0.001** |
| Food (restricted) | -0.319 | 0.444 | 0.517 | 1 | 0.472 |

| Random effect | Variance | *sd* | Number of groups |
|---|---|---|---|
| Family ID (intercept) | 0.123 | 0.351 | 26 |

Conditional part

| Fixed effect | Estimate | *SE* | $\chi^2$ | *df* | *P* |
|---|---|---|---|---|---|
| Intercept (24°C, control) | 1.082 | 0.044 | | | |
| Temperature (28°C) | -0.058 | 0.054 | 1.150 | 1 | 0.284 |
| Food (restricted) | -0.032 | 0.054 | 0.349 | 1 | 0.555 |

| Random effect | Variance | *sd* | Number of groups |
|---|---|---|---|
| Family ID (intercept) | <0.001 | <0.001 | 26 |

(iii) Exclusion of the non-significant interactions did not significantly reduce model fit in the final model

| | *df* | AIC | BIC | Log-likelihood | Deviance | $\chi^2_2$ | *P* |
|---|---|---|---|---|---|---|---|
| Initial model (i) | 9 | 706.4 | 736.8 | -344.19 | 688.39 | | |
| Final model (ii) | 11 | 709.3 | 746.4 | -343.63 | 687.25 | 1.134 | 0.567 |

**(B) Total offspring number**

(i) Initial model with the interactive effect

Zero-inflation part

| Fixed effect | Estimate | *SE* | $\chi^2$ | *df* | *P* |
|---|---|---|---|---|---|
| Intercept (24°C, control) | -2.967 | 0.630 | 22.168 | 1 | **<0.001** |
| Temperature (28°C) | 1.734 | 0.692 | 6.283 | 1 | **0.012** |
| Food (restricted) | -0.349 | 0.959 | 0.133 | 1 | 0.716 |
| Temperature (28°C) * Food (restricted) | 0.009 | 1.081 | <0.001 | 1 | 0.993 |

| Random effect | Variance | *sd* | Number of groups |
|---|---|---|---|
| Family ID (intercept) | 0.114 | 0.338 | 26 |

Conditional part

| Fixed effect | Estimate | *SE* | $\chi^2$ | *df* | *P* |
|---|---|---|---|---|---|
| Intercept (24°C, control) | 3.552 | 0.070 | 2571.453 | 1 | **<0.001** |
| Temperature (28°C) | -0.445 | 0.108 | 17.115 | 1 | **<0.001** |
| Food (restricted) | 0.061 | 0.090 | 0.463 | 1 | 0.496 |
| Temperature (28°C) * Food (restricted) | -0.323 | 0.158 | 4.179 | 1 | **0.041** |

| Random effect | Variance | *sd* | Number of groups |
|---|---|---|---|
| Family ID (intercept) | 0.019 | 0.138 | 26 |

(ii) Final model excluding the non-significant interaction

Zero-inflation part

| Fixed effect | Estimate | *SE* | $\chi^2$ | *df* | *P* |
|---|---|---|---|---|---|
| Intercept (24°C, control) | -2.970 | 0.527 | | | |
| Temperature (28°C) | 1.738 | 0.532 | 10.687 | 1 | **0.001** |
| Food (restricted) | -0.342 | 0.449 | 0.579 | 1 | 0.447 |

| Random effect | Variance | *sd* | Number of groups |
|---|---|---|---|
| Family ID (intercept) | 0.114 | 0.338 | 26 |

Conditional part

| Fixed effect | Estimate | *SE* | $\chi^2$ | *df* | *P* |
|---|---|---|---|---|---|
| Intercept (24°C, control) | 3.552 | 0.070 | 2571.440 | 1 | **<0.001** |
| Temperature (28°C) | -0.445 | 0.108 | 17.115 | 1 | **<0.001** |
| Food (restricted) | 0.061 | 0.090 | 0.464 | 1 | 0.496 |
| Temperature (28°C)*Food (restricted) | -0.323 | 0.158 | 4.181 | 1 | **0.041** |

| Random effect | Variance | *sd* | Number of groups |
|---|---|---|---|
| Family ID (intercept) | 0.019 | 0.138 | 26 |

Given a significant interaction in the conditional part, we ran the pairwise comparison:

| Temperature | contrast | ratio | SE | *df* | t.ratio | *P* |
|---|---|---|---|---|---|---|
| 28°C | control vs restricted diet | 1.30 | 0.168 | 206 | 2.026 | **0.044** |
| 24°C | control vs restricted diet | 0.94 | 0.085 | 206 | -0.681 | 0.497 |

(iii) Exclusion of the non-significant interactions did not significantly reduce model fit in the final model

| | *df* | AIC | BIC | Loglikelihood | Deviance | $\chi^2_1$ | *P* |
|---|---|---|---|---|---|---|---|
| Initial model (i) | 10 | 1681.4 | 1715.2 | -830.71 | 1661.4 | <0.001 | 0.993 |
| Final model (ii) | 11 | 1683.4 | 1720.5 | -830.71 | 1661.4 | | |

### (C) Total egg number

(i) Initial model with the interactive effect

| Fixed effect | Estimate | *SE* | $\chi^2$ | *df* | *P* |
|---|---|---|---|---|---|
| Intercept (24°C, control) | 16.226 | 0.686 | 559.241 | 1 | **<0.001** |
| Temperature (28°C) | -8.991 | 0.979 | 84.277 | 1 | **<0.001** |
| Food (restricted) | -0.048 | 0.834 | 0.003 | 1 | 0.954 |
| Temperature (28°C) * Food (restricted) | 1.375 | 1.391 | 0.977 | 1 | 0.323 |

| Random effect | Variance | *sd* | Number of groups |
|---|---|---|---|
| Family ID (intercept) | 3.081 | 1.755 | 26 |
| Residuals | 17.929 | 4.234 | |

(ii) Final model excluding the non-significant interaction

| Fixed effect | Estimate | *SE* | $\chi^2$ | *df* | *P* |
|---|---|---|---|---|---|
| Intercept (24°C, control) | 15.997 | 0.642 | | | |
| Temperature (28°C) | -8.302 | 0.691 | 144.457 | 1 | **<0.001** |
| Food (restricted) | 0.449 | 0.668 | 0.451 | 1 | 0.502 |

| Random effect | Variance | *sd* | Number of groups |
|---|---|---|---|
| Family ID (intercept) | 2.901 | 1.703 | 26 |
| Residuals | 18.124 | 4.257 | |

(iii) Exclusion of the non-significant interactions did not significantly reduce model fit in the final model

| | *df* | AIC | BIC | Loglikelihood | Deviance | $\chi^2_1$ | *P* |
|---|---|---|---|---|---|---|---|
| Final model (ii) | 5 | 1008.2 | 1023.9 | -499.09 | 998.17 | | |
| Initial model (i) | 6 | 1009.2 | 1028.0 | -498.6 | 997.2 | 0.971 | 0.325 |

(iv) An additional model testing the effect of absolute body length

| Fixed effect | Estimate | *SE* | $\chi^2$ | *df* | *P* |
|---|---|---|---|---|---|
| Intercept (24°C, control) | 15.574 | 0.628 | | | |
| Temperature (28°C) | -7.471 | 0.708 | 111.447 | 1 | **<0.001** |
| Food (restricted) | 0.598 | 0.647 | 0.854 | 1 | 0.356 |
| Week 18 body size (standardized) | 1.344 | 0.380 | 12.482 | 1 | **<0.001** |

| Random effect | Variance | *sd* | Number of groups |
|---|---|---|---|
| Family ID (intercept) | 2.591 | 1.610 | 26 |
| Residuals | 16.944 | 4.116 | |

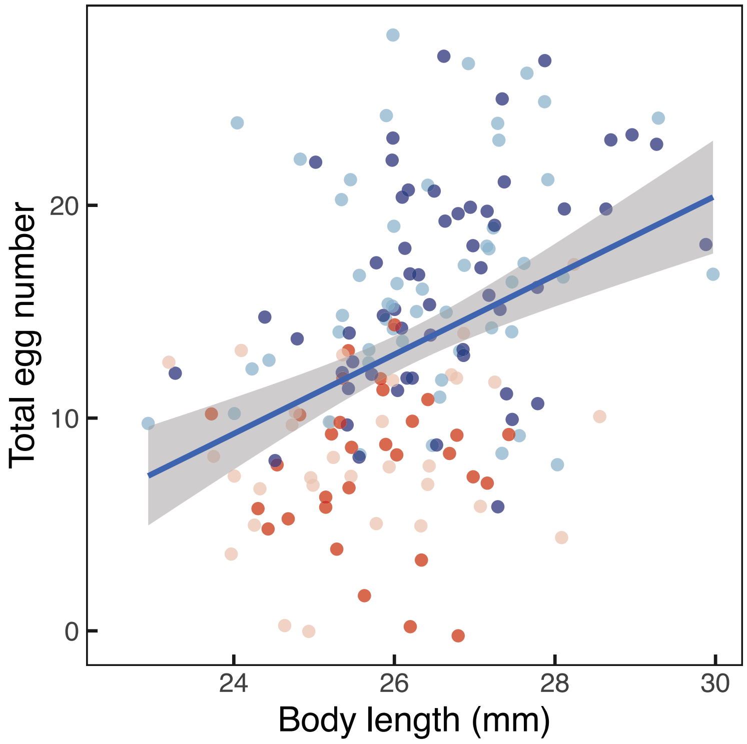


**Figure S1. Relationship between female size and total egg number.** Dot color indicates the treatments: dark blue = control temperature-control diet; light blue = control temperature-restricted diet, dark red = warm temperature-control diet; light red = warm temperature-restricted diet. Body size-dependence of total egg number across all females is shown using a regression line with the 95% confidence interval.

### (D) Embryo number

(i) Initial model with the interactive effect

Zero-inflation part

| Fixed effect | Estimate | *SE* | $\chi^2$ | *df* | *P* |
|---|---|---|---|---|---|
| Intercept (24°C, control) | -1.106 | 0.315 | 12.332 | 1 | **<0.001** |
| Temperature (28°C) | 0.852 | 0.496 | 2.957 | 1 | 0.086 |
| Food (restricted) | 0.192 | 0.439 | 0.192 | 1 | 0.661 |
| Temperature (28°C) * Food (restricted) | -0.171 | 0.695 | 0.060 | 1 | 0.806 |

| Random effect | Variance | *sd* | Number of groups |
|---|---|---|---|
| Family ID (intercept) | 0.035 | 0.188 | 26 |

Conditional part

| Fixed effect | Estimate | *SE* | $\chi^2$ | *df* | *P* |
|---|---|---|---|---|---|
| Intercept (24°C, control) | 2.693 | 0.068 | 1576.891 | 1 | **<0.001** |
| Temperature (28°C) | -0.879 | 0.166 | 27.940 | 1 | **<0.001** |
| Food (restricted) | -0.035 | 0.095 | 0.134 | 1 | 0.714 |
| Temperature (28°C) * Food (restricted) | -0.123 | 0.244 | 0.252 | 1 | 0.616 |

| Random effect | Variance | *sd* | Number of groups |
|---|---|---|---|
| Family ID (intercept) | 0.007 | 0.082 | 26 |

(ii) Final model excluding the non-significant interaction

Zero-inflation part

| Fixed effect | Estimate | *SE* | $\chi^2$ | *df* | *P* |
|---|---|---|---|---|---|
| Intercept (24°C, control) | -1.076 | 0.280 | | | |
| Temperature (28°C) | 0.767 | 0.349 | 4.821 | 1 | **0.028** |
| Food (restricted) | 0.134 | 0.341 | 0.154 | 1 | 0.695 |

| Random effect | Variance | *sd* | Number of groups |
|---|---|---|---|
| Family ID (intercept) | 0.031 | 0.176 | 26 |

Conditional part

| Fixed effect | Estimate | *SE* | $\chi^2$ | *df* | *P* |
|---|---|---|---|---|---|
| Intercept (24°C, control) | 2.701 | 0.066 | | | |
| Temperature (28°C) | -0.937 | 0.122 | 59.151 | 1 | **<0.001** |
| Food (restricted) | -0.053 | 0.088 | 0.370 | 1 | 0.543 |

| Random effect | Variance | *sd* | Number of groups |
|---|---|---|---|
| Family ID (intercept) | 0.007 | 0.084 | 26 |

(iii) Exclusion of the non-significant interactions did not significantly reduce model fit in the final model

| | *df* | AIC | BIC | Loglikelihood | Deviance | $\chi^2_2$ | *P* |
|---|---|---|---|---|---|---|---|
| Final model (ii) | 9 | 913.07 | 941.34 | -447.53 | 895.07 | | |
| Initial model (i) | 11 | 916.77 | 951.33 | -447.39 | 894.77 | 0.297 | 0.862 |

(iv) An additional model testing the effect of absolute body length

Zero-inflation part

| Fixed effect | Estimate | *SE* | $\chi^2$ | *df* | *P* |
|---|---|---|---|---|---|
| Intercept (24°C, control) | -1.259 | 0.309 | | | |
| Temperature (28°C) | 1.113 | 0.393 | 8.008 | 1 | **0.005** |
| Food (restricted) | 0.176 | 0.348 | 0.255 | 1 | 0.613 |
| Week 18 body size (standardized) | 0.461 | 0.204 | 5.085 | 1 | **0.024** |

| Random effect | Variance | *sd* | Number of groups |
|---|---|---|---|
| Family ID (intercept) | 0.077 | 0.277 | 26 |

Conditional part

| Fixed effect | Estimate | *SE* | $\chi^2$ | *df* | *P* |
|---|---|---|---|---|---|
| Intercept (24°C, control) | 2.660 | 0.069 | | | |
| Temperature (28°C) | -0.855 | 0.124 | 47.514 | 1 | **<0.001** |
| Food (restricted) | -0.025 | 0.086 | 0.087 | 1 | 0.768 |
| Week 18 body size (standardized) | 0.118 | 0.052 | 5.049 | 1 | **0.025** |

| Random effect | Variance | *sd* | Number of groups |
|---|---|---|---|
| Family ID (intercept) | 0.012 | 0.112 | 26 |

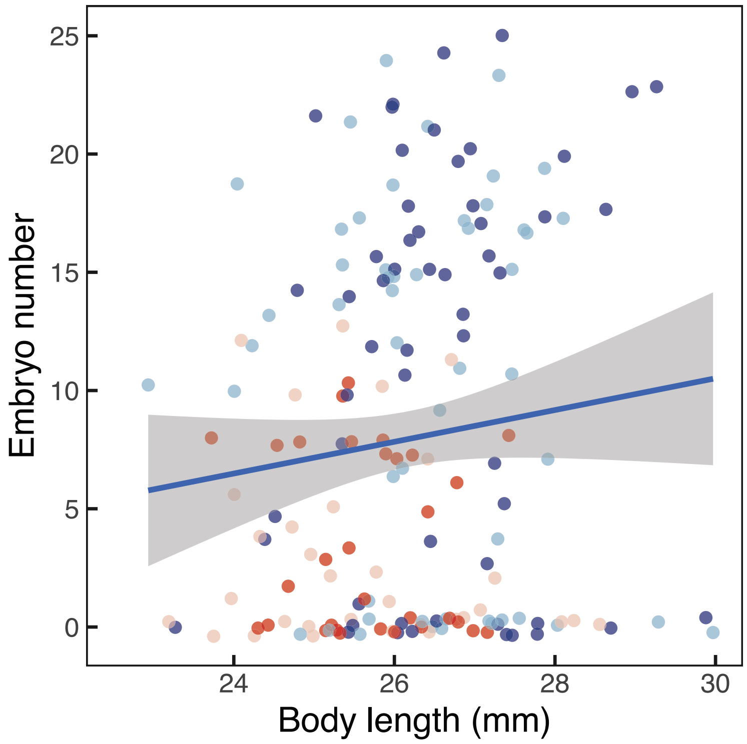


**Figure S2. Relationship between female size and embryo number.** Dot color indicates the treatments: dark blue = control temperature-control diet; light blue = control temperature-restricted diet, dark red = warm temperature-control diet; light red = warm temperature-restricted diet. Body size-dependence of embryo number across all females is shown using a regression line with the 95% confidence interval.

**Table S5. effects of temperature and early-life food availability on mortality and somatic investment**

**(A) Mortality**

(i) Comparison of survival curves among four temperature-diet treatments

| Treatment group | *N* | Observed | Expected | (Observed – Expected)^2/Expected |
|---|---|---|---|---|
| 24°C-control diet | 57 | 1 | 11.3 | 9.35 |
| 24°C-restricted diet | 52 | 0 | 11.1 | 11.10 |
| 28°C-control diet | 55 | 23 | 11.5 | 11.46 |
| 28°C-restricted diet | 52 | 20 | 10.1 | 9.64 |

(ii) Comparison of survival curves between two temperatures

| Treatment group | *N* | Observed | Expected | (Observed – Expected)^2/Expected |
|---|---|---|---|---|
| 24°C | 109 | 1 | 22.4 | 20.4 |
| 28°C | 107 | 43 | 21.6 | 21.1 |

(iii) Comparison of survival curves between two diet treatments

| Treatment group | *N* | Observed | Expected | (Observed – Expected)^2/Expected |
|---|---|---|---|---|
| Control diet | 112 | 24 | 22.8 | 0.066 |
| Restricted diet | 104 | 20 | 21.2 | 0.071 |

(iv) Comparison of survival curves between two diet treatments at 24°C

| Treatment group | *N* | Observed | Expected | (Observed – Expected)^2/Expected |
|---|---|---|---|---|
| Control diet | 57 | 1 | 0.6 | 0.267 |
| Restricted diet | 52 | 0 | 0.4 | 0.400 |

(v) Comparison of survival curves between two diet treatments at 28°C

| Treatment group | *N* | Observed | Expected | (Observed – Expected)^2/Expected |
|---|---|---|---|---|
| Control diet | 55 | 23 | 23 | <0.001 |
| Restricted diet | 52 | 20 | 20 | <0.001 |

### (B) Relative telomere length

#### (i) Initial model with the interactive effect

| Fixed effect | Estimate | *SE* | $\chi^2$ | *df* | *P* |
|---|---|---|---|---|---|
| Intercept (24°C, control) | 0.971 | 0.053 | 338.630 | 1 | **<0.001** |
| Temperature (28°C) | 0.013 | 0.067 | 0.041 | 1 | 0.840 |
| Food (restricted) | 0.042 | 0.066 | 0.411 | 1 | 0.522 |
| Temperature (28°C) * Food (restricted) | 0.019 | 0.094 | 0.039 | 1 | 0.843 |

| Random effect | Variance | *sd* | Number of groups |
|---|---|---|---|
| Family ID (intercept) | 0.014 | 0.120 | 26 |
| Residuals | 0.064 | 0.254 | |

#### (ii) Final model excluding the non-significant interaction

| Fixed effect | Estimate | *SE* | $\chi^2$ | *df* | *P* |
|---|---|---|---|---|---|
| Intercept (24°C, control) | 0.966 | 0.048 | | | |
| Temperature (28°C) | 0.023 | 0.047 | 0.233 | 1 | 0.629 |
| Food (restricted) | 0.051 | 0.048 | 1.165 | 1 | 0.281 |

| Random effect | Variance | *sd* | Number of groups |
|---|---|---|---|
| Family ID (intercept) | 0.015 | 0.120 | 26 |
| Residuals | 0.064 | 0.254 | |

#### (iii) Model comparison

| | *Df* | AIC | BIC | Loglikelihood | Deviance | $\chi^2_1$ | *P* |
|---|---|---|---|---|---|---|---|
| Final model (ii) | 5 | 38.73 | 52.667 | -14.365 | 28.73 | | |
| Initial model (i) | 6 | 40.69 | 57.415 | -14.345 | 28.69 | 0.039 | 0.843 |

(iv) An additional model testing the effect of absolute body length

| Fixed effect | Estimate | *SE* | $\chi^2$ | *df* | *P* |
|---|---|---|---|---|---|
| Intercept (24°C, control) | 0.965 | 0.049 | | | |
| Temperature (28°C) | 0.025 | 0.051 | 0.243 | 1 | 0.622 |
| Food (restricted) | 0.052 | 0.048 | 1.178 | 1 | 0.278 |
| Week 18 size (standardized) | 0.003 | 0.029 | 0.014 | 1 | 0.905 |

| Random effect | Variance | *sd* | Number of groups |
|---|---|---|---|
| Family ID (intercept) | 0.014 | 0.120 | 26 |
| Residuals | 0.064 | 0.254 | |

(v) An additional model testing the effect of chronological age (i.e., birth to age of testing)

| Fixed effect | Estimate | *SE* | $\chi^2$ | *df* | *P* |
|---|---|---|---|---|---|
| Intercept (24°C, control) | 1.006 | 0.054 | | | |
| Temperature (28°C) | -0.005 | 0.051 | 0.010 | 1 | 0.920 |
| Food (restricted) | 0.005 | 0.057 | 0.008 | 1 | 0.929 |
| Absolute age (standardized) | 0.046 | 0.032 | 2.045 | 1 | 0.153 |

| Random effect | Variance | *sd* | Number of groups |
|---|---|---|---|
| Family ID (intercept) | 0.013 | 0.113 | 26 |
| Residuals | 0.064 | 0.253 | |

### (C) Immune response - size of swelling (µm)

#### (i) Initial model with the interactive effect

Zero-inflation part

| Fixed effect | Estimate | *SE* | $\chi^2$ | *df* | *P* |
|---|---|---|---|---|---|
| Intercept | -3.415 | 0.448 | 58.011 | 1 | **<0.001** |

Conditional part

| Fixed effect | Estimate | *SE* | $\chi^2$ | *df* | *P* |
|---|---|---|---|---|---|
| Intercept (24°C, control) | 4.287 | 0.107 | 1606.047 | 1 | **<0.001** |
| Pre-injection thickness (standardized) | -0.041 | 0.056 | 0.537 | 1 | 0.464 |
| Temperature (28°C) | -0.038 | 0.149 | 0.066 | 1 | 0.798 |
| Food (restricted) | 0.050 | 0.128 | 0.151 | 1 | 0.697 |
| Temperature (28°C) * Food (restricted) | -0.184 | 0.210 | 0.763 | 1 | 0.382 |

| Random effect | Variance | *sd* | Number of groups |
|---|---|---|---|
| Family ID (intercept) | 0.066 | 0.258 | 26 |

#### (ii) Final model excluding the non-significant interaction

Zero-inflation part

| Fixed effect | Estimate | *SE* | $\chi^2$ | *df* | *P* |
|---|---|---|---|---|---|
| Intercept | -3.438 | 0.456 | 56.718 | 1 | **<0.001** |

Conditional part

| Fixed effect | Estimate | *SE* | $\chi^2$ | *df* | *P* |
|---|---|---|---|---|---|
| Intercept (24°C, control) | 4.316 | 0.102 | | | |
| Pre-injection thickness (standardized) | -0.045 | 0.056 | 0.635 | 1 | 0.426 |
| Temperature (28°C) | -0.131 | 0.107 | 1.499 | 1 | 0.221 |
| Food (restricted) | -0.017 | 0.103 | 0.026 | 1 | 0.871 |

| Random effect | Variance | *sd* | Number of groups |
|---|---|---|---|
| Family ID (intercept) | 0.074 | 0.273 | 26 |

#### (iii) Model comparison

| | *df* | AIC | BIC | Loglikelihood | Deviance | $\chi^2_1$ | *P* |
|---|---|---|---|---|---|---|---|
| Final model (ii) | 7 | 1836.4 | 1858.6 | -911.2 | 1822.4 | | |
| Initial model (i) | 8 | 1837.6 | 1863.0 | -910.8 | 1821.6 | 0.765 | 0.382 |

(iv) An additional model testing the effect of absolute body length

Zero-inflation part

| Fixed effect | Estimate | *SE* | $\chi^2$ | *df* | *P* |
|---|---|---|---|---|---|
| Intercept | -3.438 | 0.457 | 56.68 | 1 | **<0.001** |

Conditional part

| Fixed effect | Estimate | *SE* | $\chi^2$ | *df* | *P* |
|---|---|---|---|---|---|
| Intercept (24°C, control) | 4.319 | 0.103 | | | |
| Pre-injection thickness (standardized) | -0.040 | 0.060 | 0.448 | 1 | 0.503 |
| Temperature (28°C) | -0.138 | 0.111 | 1.548 | 1 | 0.214 |
| Food (restricted) | -0.017 | 0.103 | 0.028 | 1 | 0.868 |
| Week 18 body size (standardized) | -0.013 | 0.057 | 0.055 | 1 | 0.815 |

| Random effect | Variance | *sd* | Number of groups |
|---|---|---|---|
| Family ID (intercept) | 0.075 | 0.275 | 26 |

### (D) Relative gut length (log-transformed)

(i) Initial model with the interactive effect

| Fixed effect | Estimate | *SE* | $\chi^2$ | *df* | *P* |
|---|---|---|---|---|---|
| Intercept (24°C, control) | 3.375 | 0.022 | 22872.070 | 1 | **<0.001** |
| Temperature (28°C) | 0.089 | 0.036 | 6.124 | 1 | **0.013** |
| Food (restricted) | 0.004 | 0.029 | 0.015 | 1 | 0.904 |
| Log-transformed Week 18 size (standardized) | 0.072 | 0.013 | 29.456 | 1 | **<0.001** |
| Temperature (28°C) * Food (restricted) | 0.044 | 0.049 | 0.821 | 1 | 0.365 |

| Random effect | Variance | *sd* | Number of groups |
|---|---|---|---|
| Family ID (intercept) | 0.001 | 0.037 | 26 |
| Residuals | 0.023 | 0.150 | |

(ii) Final model excluding the non-significant interaction

| Fixed effect | Estimate | *SE* | $\chi^2$ | *df* | *P* |
|---|---|---|---|---|---|
| Intercept (24°C, control) | 3.367 | 0.021 | | | |
| Temperature (28°C) | 0.112 | 0.026 | 18.255 | 1 | **<0.001** |
| Food (restricted) | 0.020 | 0.023 | 0.705 | 1 | 0.401 |
| Log Week 18 body size (standardized) | 0.073 | 0.013 | 29.794 | 1 | **<0.001** |

| Random effect | Variance | *sd* | Number of groups |
|---|---|---|---|
| Family ID (intercept) | 0.001 | 0.035 | 26 |
| Residuals | 0.023 | 0.151 | |

(iii) Model comparsion

| | *df* | AIC | BIC | Loglikelihood | Deviance | $\chi^2_1$ | *P* |
|---|---|---|---|---|---|---|---|
| Final model (ii) | 6 | -141.8 | -122.9 | 76.875 | -153.75 | | |
| Initial model (i) | 7 | -140.6 | -118.6 | 77.284 | -154.57 | 0.818 | 0.336 |

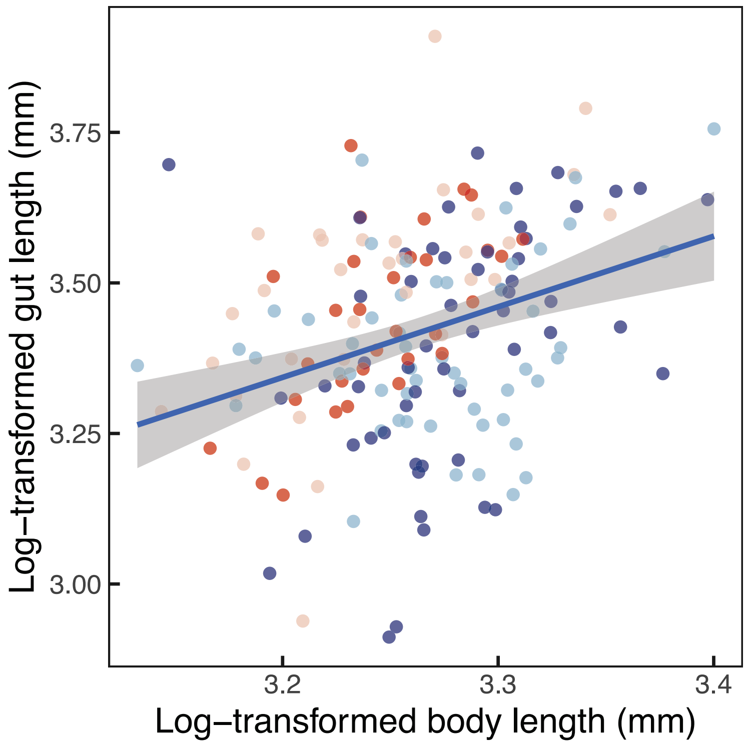


**Figure S3. Relationship between female size (log-transformed) and gut length (log-transformed).** Dot color indicates the treatments: dark blue = control temperature-control diet; light blue = control temperature-restricted diet, dark red = warm temperature-control diet; light red = warm temperature-restricted diet. The allometric relationship across all females is shown using a regression line with the 95% confidence interval.

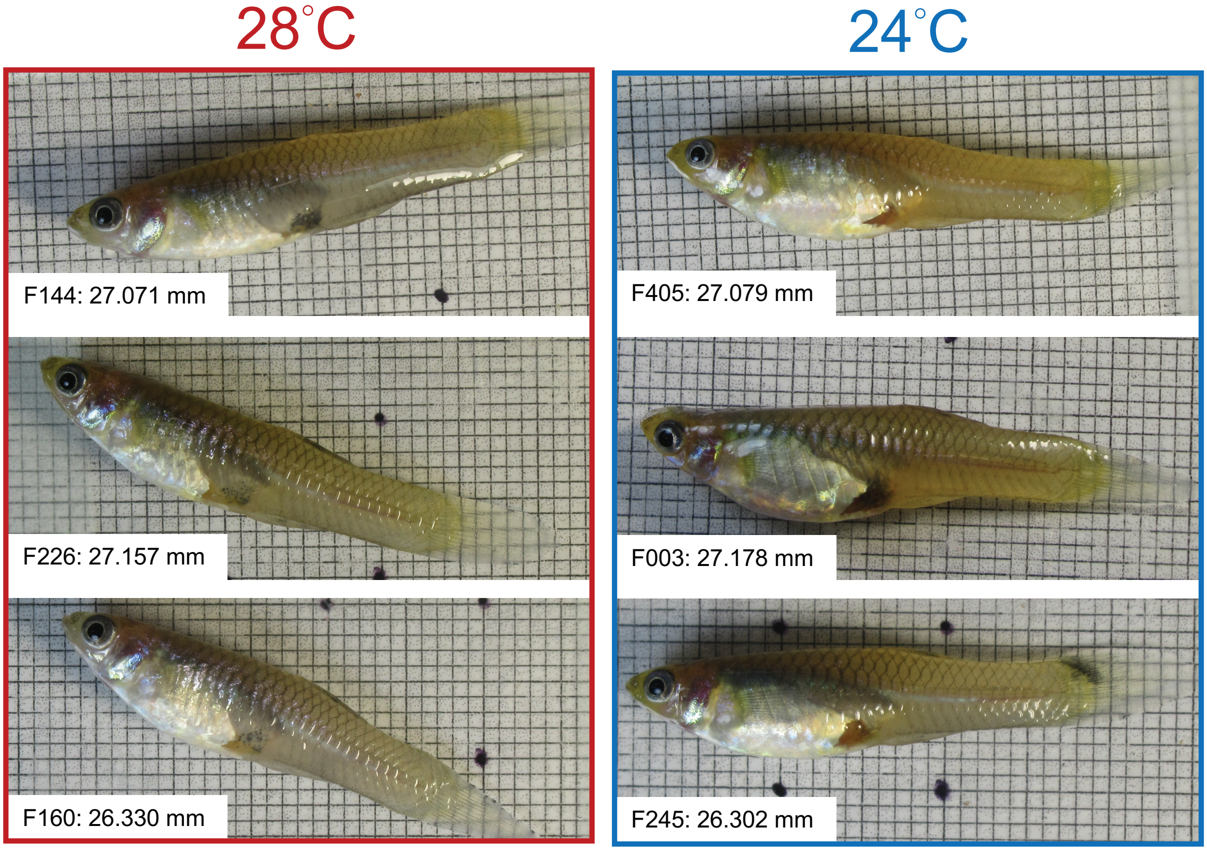


**Figure S4. Photographs of size-matched females at 28°C (left; red frame) versus 24°C (right; blue frame) at week 18.** Female identity and body length (mm) were shown. Each background square is 1 × 1 mm².
